# Spatially explicit ecological modeling improves empirical characterization of dispersal

**DOI:** 10.1101/789156

**Authors:** Petteri Karisto, Frédéric Suffert, Alexey Mikaberidze

## Abstract

Dispersal is a key ecological process, but remains difficult to measure. By recording numbers of dispersed individuals at different distances from the source one can acquire a dispersal gradient. Although dispersal gradients contain information on dispersal, they are influenced by the spatial extent of the source. How can we separate the two contributions to extract knowledge on dispersal?

One could use a small, point-like source for which a dispersal gradient represents a dispersal kernel, which quantifies the probability of an individual dispersal event from a source to a destination point. However, the validity of this approximation cannot be established before conducting measurements.

We formulated a theory that incorporates the spatial extent of sources to estimate dispersal kernels from dispersal gradients. We re-analyzed published dispersal gradients for three major plant pathogens. We also demonstrated using simulations that this approach provides more accurate estimates of dispersal kernels across biologically plausible scenarios. We concluded that the three plant pathogens disperse over substantially shorter distances compared to conventional estimates. Using this method, a significant proportion of published dispersal gradients can be re-analyzed to improve our knowledge about spatial scales of dispersal. Thus, our results can boost progress in characterization of dispersal across taxa.

## Introduction

Dispersal is an important component of many life histories, because individual organisms commonly move from one location to another in order to survive and reproduce. Empirical characterization of dispersal has been a major theme in ecological research for over a hundred years (for example Heald, 1913; Bullock et al., 2017) and dispersal measurements are often supported by solid theoretical frameworks (Nathan et al., 2012). In spite of that, we still have far fewer datasets describing, for example, plant dispersal than plant demography, because dispersal remains difficult to measure (Bullock et al., 2017). We identified and resolved one of the key challenges that hinders progress in empirical characterization of dispersal: we incorporated the spatial extent of dispersal sources in the analysis of dispersal measurements.

One approach to measure dispersal is to use a spatially localized source of dispersing individuals and record the number of dispersed individuals as a function of the distance from this source. A localized source is expected to produce a dispersal gradient: on average, more individuals will travel only short distances from the source, while fewer individuals will travel further, leading to a decreasing trend over distance. Many studies of dispersal report measured dispersal gradients. Although they do contain information on relevant dispersal properties of the population under study, they are also strongly influenced by the spatial extent of the source (Zadoks and Schein, 1979; Ferrandino, 1996; Cousens and Rawlinson, 2001).

How can we separate the two contributions to extract more general knowledge about dispersal from specific dispersal gradients? This can be achieved using a sound mathematical description of dispersal with the help of dispersal kernels. A dispersal kernel quantifies the probability of an individual dispersal event from a source point to a destination point, and thereby contains comprehensive knowledge about dispersal properties. Technically, a dispersal kernel is a probability density function that depends on the spatial location of the destination point (“dispersal location kernel”, Nathan et al., 2012). To characterize dispersal, we need to estimate dispersal kernels based on observed dispersal gradients. If dispersal measurements could be done using an infinitely small source, i.e., a point source, then observed dispersal gradients would correspond to dispersal kernels. In practice, however, sources need to have a certain area to yield any dispersing propagules, and usually the area should be considerably large to secure an observable number of dispersal events to measure dispersal gradients.

To estimate dispersal kernels, one could use source areas that are small enough to be considered as points (the point source approximation). But how small do the source areas need to be? A rule of thumb that received a great deal of attention in the literature states that a point source should have “a diameter smaller than 1% of the gradient length” (Zadoks and Schein, 1979). This rule of thumb is misleading: To determine whether the source is small enough so that the dispersal gradient captures the shape of the dispersal kernel, the size of the source should be compared with the characteristic distance of dispersal (i.e., the distance over which the dispersal kernel changes substantially), rather than the gradient length (i.e., the extent of measurements). This represents a challenge for the design of dispersal experiments that aim to achieve a point-like source, because we do not know the characteristic distance of dispersal before doing the measurements. Therefore, we cannot establish sound criteria for the validity of the point source approximation in advance of conducting measurements.

As a result of this uncertainty, “point” sources of various sizes appear in the literature: an adult tree (lichen *Lobaria pulmonaria,* Werth et al., 2006), circles of 1.6m diameter (thistles *Carduus nutans, Carduus acanthoides,* Skarpaas and Shea, 2007), circles of 0.5 m diameter (garlic mustard *Alliaria petiolata,* Loebach and Anderson, 2018), 4m^2^ squares (wine raspberry *Rubus phoenicolasius,* Japanese barberry *Berberis thunbergii,* multiflora rose *Rosa multiflora,* and Japanese stiltgrass *Microstegium vimineum,* Emsweller et al., 2018), routes of individual sampling dives (coral reef fish *Elacatinus lori,* D’Aloia et al., 2015), and even entire agricultural fields (oilseed rape *Brassica napus,* Devaux et al., 2006). These studies reported valuable dispersal gradients, but it is unlikely that the sources used were sufficiently small to be considered as point sources in all cases. Hence, using dispersal gradients measured in these studies as proxies for dispersal kernels may be unjustified. Spatially explicit modeling has been suggested (Greene and Calogeropoulos, 2002) to address this problem and was used in some modeling studies (Clark et al., 1999; Shaw et al., 2006), but it was not widely adopted in the literature on dispersal measurements.

In this study, we devised a systematic approach to estimate dispersal kernels from dispersal gradients without using the point source approximation. For this purpose, we combined theory, analysis of empirical data and numerical simulations. We first formulated a theory that incorporates dispersal from a spatially extended source considering each point within the source area as an independent point source (the spatially explicit approach). We highlighted how seemingly abstract mathematical properties of widely used kernel functions (exponential, Gaussian and power-law, Nathan et al., 2012) can inform experimental design. Then, we re-analyzed published empirical datasets on three major plant pathogens with contrasting spatial scales of dispersal and conducted comprehensive numerical simulations. In this way, we demonstrated how this approach allows to estimate dispersal kernels more accurately than using the point source approximation.

## Theory

The probability of dispersal from a source point *p_s_* = (*x_s_,y_s_*) to a destination point *p_d_* = (*x_d_,y_d_*) is given by the *dispersal location kernel* (hereafter “dispersal kernel”). It is typically a monotonically decreasing function of the distance between the points 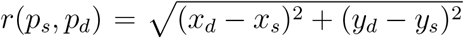. The spatial scale of dispersal is characterized by the mean dispersal distance, 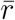, which can be calculated from the dispersal kernel.

To estimate a dispersal kernel using a dispersal gradient produced by an area source, we consider the cumulative effect of all point-to-point dispersal events from the source to the destination. This is achieved by taking a sum over the individual points comprising the source to calculate their combined contribution to the dispersed population at a certain destination point (as in Shaw et al., 2006, Eq. (4.6)). Similarly, the sum over all points of the destination area gives the total number of individuals that moved there from the source:

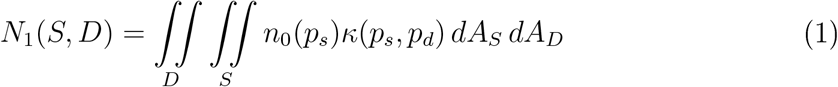

where *S* = {*p_s_*} is the source area, *D* = {*p_d_*} is the destination area, *n*_0_(*p_s_*) is the density of individuals within *S* before dispersal and *κ*(*p_s_,p_d_*) is the dispersal kernel. The area integrals in Eq. (1) sum up the contributions of all source points in *S* to all destination points in *D* to compute the total population in *D*. Eq. (1) provides a valid description of the dispersal process when the overall population size is sufficiently large so that stochastic fluctuations in the numbers of dispersed individuals can be neglected. When the populations before dispersal (*n*_0_(*p_s_*)) and after dispersal (*N*_1_) are measured, the only unknown in Eq. (1) remains the dispersal kernel. Given a specific kernel function, its parameters can be estimated by fitting this function to the observed data. Eq. (1) offers a way to estimate dispersal kernel parameters that takes into account the spatial extent of both the source and the destination.

A simpler approach that is much more common is to fit a function of one spatial coordinate *x* to dispersal gradient data. For example, the function

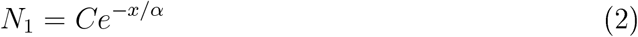

can be fitted to dispersal gradient data to estimate the scale parameter *α* of the expo-nential kernel in Box 1 (for example, Saint-Jean et al., 2004). If both the source and the destination can be considered as points, the above approach provides a correct estimate of *α*, because the function in Eq. (2) is the same as the exponential dispersal kernel [Eq. (3) in Box 1] up to a constant factor. This approach works for any kernel function if both the source and the destination can be considered as points.

#### Box 1. Dispersal kernels

##### Exponential kernel

is defined as

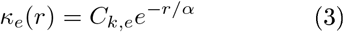

where *α* is the scale parameter, *k* ∈ {1, 2} is the number of dimensions, *r* = *r*(*p_s_,p_d_*) is the Euclidean distance from the source point *p_s_* = (*x_s_,y_s_*) to the destination point *p_d_* = (*x_d_,y_d_*) (in one dimension *y_s_* = *y_d_* = 0), and *C_k,e_* is the normalization factor: *C*_1,*e*_ = 1/(2*α*), *C*_2,*e*_ = 1/(2*πα*^2^). The mean dispersal distance at *k* = 2 is 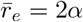.

##### Gaussian kernel

is defined as

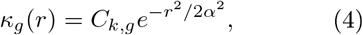

where *α* is the scale parameter, 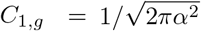, and *C*_2,*g*_ = 1/(2*πα*^2^). The mean dispersal distance at *k* = 2 is 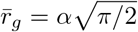.

##### Power-law kernel

is defined here as

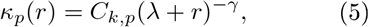

where *γ* is the shape parameter, λ is the scale parameter, *C*_1,*p*_ = (*γ* – 1)λ^*γ*–1^, *C*_2,*p*_ = (*γ* – 2)(*γ* – 1)λ^*γ*–2^/(2*π*).

The mean dispersal distance at *k* = 2 is 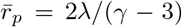 for *γ* > 3 and 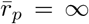 for *γ* ≤ 3.

However, when the source is extended in space, the above approach may lead to inac-curate estimates of kernel parameters, because extended sources modify dispersal gradients compared to point sources. Figure 1 illustrates such modifications for exponential, Gaussian and power-law kernels (defined in Box 1). Compare, for example, the gradients produced by the point source (source 1 in Fig. 1A) and the area source (source 4 in Fig. 1A). Extension of the source leads to a “flattening” of the gradient for the exponential and the power-law kernels, but for the Gaussian kernel it leads to a “steepening” of the gradient (cf. the dashed green curve with the dashed blue curve in Fig. 1B,D). Some studies postulated that gradients produced by spatially extended sources are “flatter” than gradients resulting from more localized sources (Zadoks and Schein, 1979; Ferrandino, 1996; Cousens and Rawlinson, 2001; Greene and Calogeropoulos, 2002). Here, we demonstrated that whether the extension of the source leads to a “flattening” or to a “steepening” of the gradient depends on the underlying kernel function. Thus, using a dispersal gradient from an extended source as a proxy for a dispersal kernel can lead to either an overestimation or an underestimation of the associated kernel parameters.

**Figure 1:**
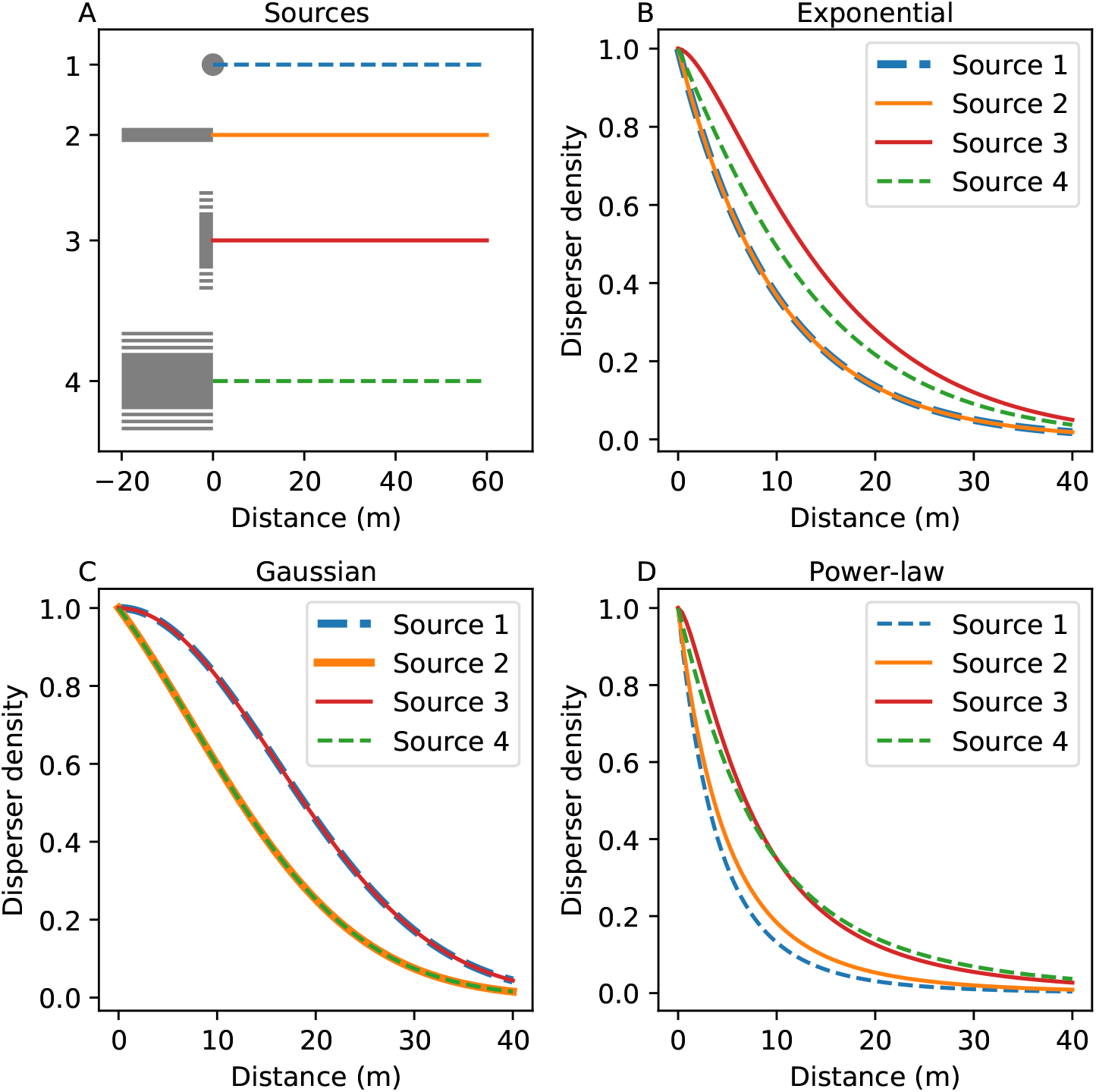
Different extensions of the source (panel A) lead to different effects on the dispersal gradients (B, C, D) depending on the dispersal kernel. Kernel parameters are chosen so that the mean dispersal distance 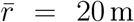 in all cases. All gradients are normalized to begin at one. (A) Four different sources (grey shapes): 1) a point source; 2) a line source, parallel to the gradient (colored lines) *x_S_* ∈ [−20m,0m]; 3) a line source perpendicular to the gradient, *y_s_* ∈ [– 100m,100 m]; 4) a rectangular area source, (*x_s_,y_s_*) ∈ [–20m,0m] × [–100m,100m], (B) With the exponential kernel (*α* = 10 m), sources 1 and 2 result in identical gradients. (C) With the Gaussian kernel (*α* = 16 m) the gradients are identical between sources 1 and 3 and between sources 2 and 4. (D) With the power-law kernel (γ = 5, λ = 20 m) all gradients are different.

Only in certain special cases the shape of the dispersal gradient does match to the shape of the dispersal kernel even when the source is extended, whereby the analysis can be simplified. If the source is extended in the direction of the measured gradient (i.e., along the *x*-axis as in source 2 in Fig. 1), and dispersal is governed by the exponential kernel, Eq. (2) will still give a correct estimate of *α*, because exponential kernels are “memoryless” (Box 2). This property allows to sum up all point sources within the source area along the *x*-axis to an equivalent virtual point source at *x* = 0. Hence, the extension of the source along the *x*-axis will not distort the dispersal gradient: in Fig. 1B the point source (source 1) and the source extended along the *x*-axis (source 2) result in same gradients (up to a constant factor). However, this is not true for Gaussian and power-law kernels: in Fig. 1C,D, source 1 and source 2 produce different gradients.

#### Box 2. Special properties of kernels.

##### Memorylessness

Exponential kernels are memoryless: when we set any point in the distribution as a starting point, the tail of the distribution will have the same shape as the entire distribution (the “past” does not affect the ‘future” probabilities; see (Ahmad and Alwasel, 1999) for a mathematical definition).

##### Separability

A function is called separable when it can be expressed as a product of other functions that depend on only one independent variable each: the variables can be separated from each other, e.g., *f*(*x, y*) = *f_x_*(*x*)*f_y_*(*y*). (Separable functions are often considered in connection with separable differential equations (Ahmad and Ambrosetti, 2015)). When a dispersal kernel is separable, the shape of the kernel along the *x*-axis does not depend on the *y*-coordinate.

If the source is extended along the *y*-axis, perpendicular to the direction of measured gradient, a similar simplification is possible for the Gaussian kernel (Fig. 1C). This kernel is separable (Box 2), which means that each point source within the line source 3 in Fig. 1A results in the same gradient along the *x*-axis. Hence, when measuring the dispersal gradient along the *x*-axis, the extension of the source along the *y*-axis does not affect the gradient and therefore does not distort the estimate of the scale parameter *a.* This property is illustrated in Fig. 1C for the Gaussian kernel: the source extended along the *y*-axis (source 3) produces the same gradient as the point source (source 1). This holds for any separable kernel, but does not hold for non-separable kernels such as exponential or power-law kernels (Fig. 1B, D). Analogous simplifications can be made when considering spatially extended destinations.

Insights presented above inform design and analysis of dispersal experiments. Gaussian and exponential kernels have been used in a number of studies to describe dispersal across a range of taxonomic groups (Table 15.1 in Nathan et al., 2012). When dispersal is governed by a memoryless (exponential) or a separable (e.g., Gaussian) kernel, appropriate line sources should be used to boost the power of the source, while maintaining the validity of the point source approximation to simplify the analysis. However, in most cases dispersal is better described by kernels that are neither memoryless nor separable (Nathan et al., 2012), such as the power-law kernel in Eq. (5). In these cases, or when the kernel function is not known before conducting measurements, dispersal gradients should be analyzed using a spatially explicit approach based on Eq. (1), as we demonstrate next.

## Analysis of empirical data

We re-analyzed published empirical data on dispersal gradients using the spatially explicit method that incorporates the spatial extent of the source and compared the outcomes with those based on the conventional point source approximation. We considered three datasets collected in field experiments investigating dispersal of major pathogens of crop plants: (i) the fungus *Zymoseptoria tritici* that causes septoria tritici blotch in wheat (Karisto et al., 2021); (ii) the fungus *Puccinia striiformis* that causes stripe (yellow) rust in wheat (Sackett and Mundt, 2005; Cowger et al., 2005); and (iii) the oomycete *Phytophthora infestans* that causes late blight in potatoes (Gregory, 1968). The three pathosystems represent contrasting modes and spatial scales of dispersal. Asexual spores of *Z. tritici* (pycnidiospores) move pre-dominantly via rain splash, while asexual spores of *P. striiformis* (urediniospores) and propagules of *P. infestans* (sporangia) are mainly wind-dispersed. Spatial scales of the experiments varied from 100 cm to 100 m. Design of experimental plots and measurements is shown in Fig. 2.

**Figure 2:**
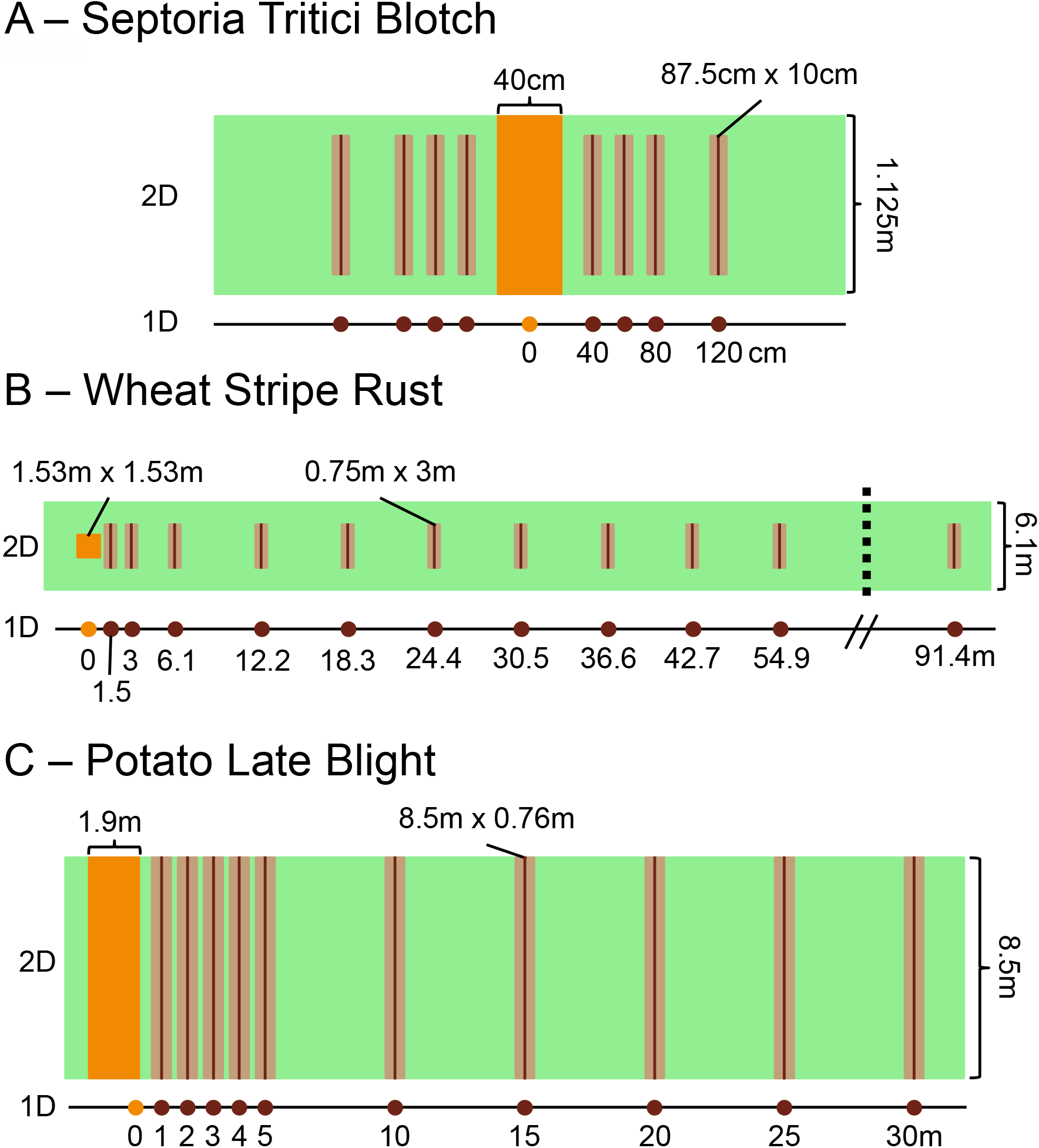
Designs of the experimental plots. (A) Septoria tritici blotch experiment (Karisto et al., 2021); (B) yellow rust experiment (Sackett and Mundt, 2005; Cowger et al., 2005); (C) potato late blight experiment (Gregory, 1968). The two-dimensional (2D) view corresponds to the spatially explicit approach; the one-dimensional (1D) view corresponds to the point source approximation. Inoculation areas are shown in orange, measurement lines in light brown and the approximated thin lines in dark brown.

In each experiment, pathogen spores were inoculated across inoculation areas within experimental plots to create area sources of dispersing populations (orange areas in Fig. 2). Then, disease gradients (disease intensity versus distance from the source) were recorded outside the inoculation areas. These gradients are called primary, because they resulted from a single cycle of pathogen reproduction. The cycle includes both spore dispersal and infection success, hence, the measured gradient reflected effective dispersal gradients of the pathogen population (analogous to the combination of seed dispersal and establishment, Klein et al., 2013). We assumed

In the analysis, we incorporated the spatial extent of the source areas, but neglected the spatial extent of the destination areas as it was considerably smaller than the characteristic dispersal distance. For each dataset, we chose an appropriate dispersal kernel function, derived the specific expression for dispersal gradients based on the more general Eq. (1) and estimated dispersal kernel parameters.

### Experimental design and data analysis

#### Septoria tritici blotch

We analyzed a subset of data collected in a larger experiment (Karisto et al., 2021) that describes dispersal of a specific pathogen strain (ST99CH_3D7). Asexual spores of *Z. tritici* were inoculated across 40 cm-wide areas in the middle of each plot (the orange rectangle in Fig. 2A). Disease intensity was measured as the density of the fungal fruiting bodies of *Z. tritici* (pycnidia) present within septoria tritici blotch lesions on leaves using automated digital image analysis (Stewart et al., 2016; Karisto et al., 2018). These measurements were conducted across thin rectangular areas situated at increasing distances from the source (we call these areas “measurement lines”; they are shown as light brown rectangles in Fig. 2A).

We first assumed that dispersing individuals originated from an infinitesimally small point at the center of the inoculation area at *x* = 0 (i.e., used the point source approximation; Fig. 2A, “1D”). Using the exponential kernel in Eq. (3) with *k* = 1, we computed the disease intensity after dispersal at a distance *r* = *x* from the source

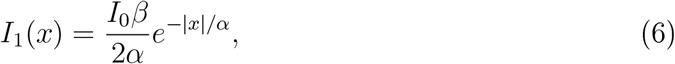

where *I*_0_ is the disease intensity at the source before dispersal and *β* is the transmission parameter comprising the probability of dispersal and the infection efficiency of fungal spores. We used the exponential kernel because it fits well to experimental data on splash-dispersed plant pathogens (Fitt et al., 1987).

Next, we lifted the point source approximation and used the spatially explicit approach. We computed the expected disease intensity after dispersal at a destination point by substituting the kernel Eq. (3) with *k* = 2 into Eq. (1) and specifying the integrals in Eq. (1) according to the plot design

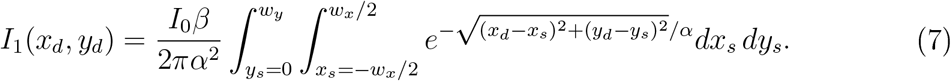

###### Box 3. Key variables and parameters.

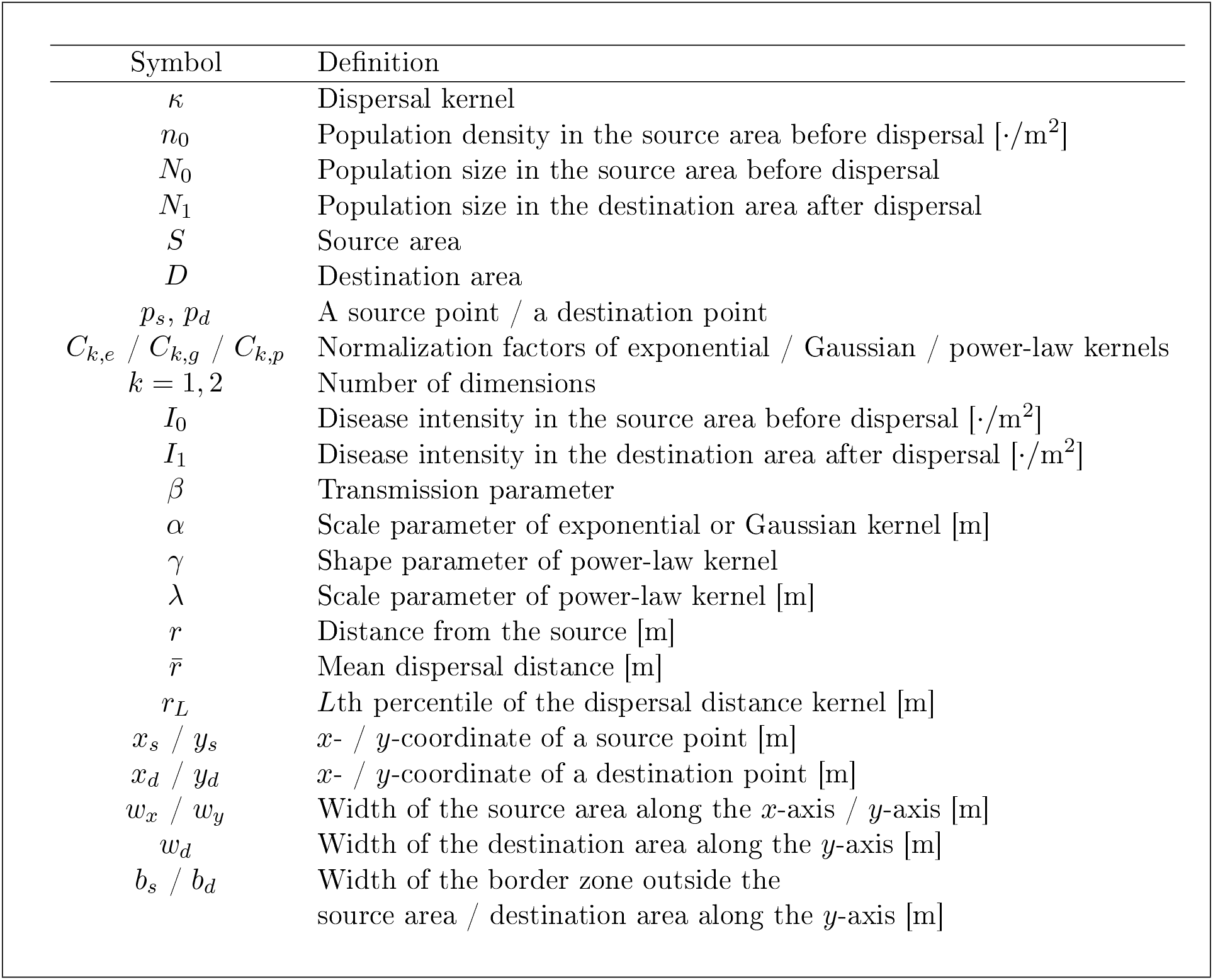

Here, we integrated across the inoculation area, (*w_x_* = 0.4 m × *w_y_* = 1.125 m), and in this way incorporated the contribution of each source point (*x_s_, y_s_*) to disease intensity at the destination point (*x_d_, y_d_*).

We further adjusted Eq. (7) to incorporate two specific features of the experimental design: (i) values recorded at different distances from the source were averages over: multiple values acquired within measurement lines (light brown rectangles in Fig. 2A) and (ii) these areas did not span the whole width of the plot but excluded its borders. With this in mind, the average disease intensity in a measurement line at a distance *x_d_* from the center of the source reads

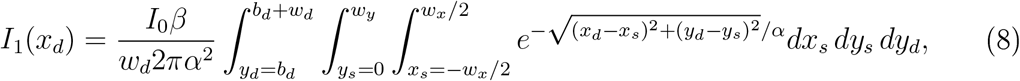

where there is an additional integration over the extent of measurement lines *y_d_* along the *y*-axis. The right-hand side of Eq. (8) is divided by the length, *w_d_*, of the measurement line (wd = *w_y_* – 2*b_d_* = 0.875 m, where *b_d_* = 0.125 m is the width of the border excluded from measurements at each end) to provide the average disease intensity across the line. In the integrals in Eq. (8), we set *x_s_* = 0 at the center of the inoculation area and *y_s_* = 0, *y_d_* = 0 at the edge of the plot.

We fitted the one-dimensional model Eq. (6) and the two-dimensional model Eq. (8) to observed dispersal gradients to estimate the scale parameter *α*.

#### Stripe rust

We analyzed a subset of data that corresponds to Hermiston 2002 and Madras 2002 trials (Sackett and Mundt, 2005; Cowger et al., 2005). Asexual spores of *P. striiformis* (urediniospores) were inoculated across 1.53 m x 1.53 m squares (the orange square in Fig. 2B) within 6.1 m-wide plots that were at least 100 m-long in the downwind direction (Cowger et al., 2005). Disease severity was measured visually as the percentage of leaf area covered by lesions (“disease severity” is a specific form of the more general term “disease intensity”, Madden et al., 2007) at increasing distances from the center of the inoculation area (from 1.5 m up to 91.4m; Fig. 2B). Disease severity was assessed as the average over O.75m-wide and 3m-long measurement lines across the middle of the plot (light brown rectangles in Fig. 2B).

We used the modified power-law kernel

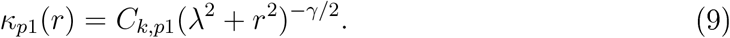

as defined by Eq. (10) of Mikaberidze et al. (2016), because it describes disease gradients of stripe rust better than exponential or Gaussian kernels. Here, r is the distance between the source point and the destination point; is the normalization factor, *k* = 1, 2 is the number of dimensions. At *k* = 1, 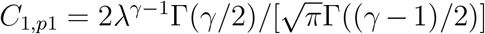, where Γ(·) is the gamma-function; at *k* = 2 *C*_2,*p*1_ = (γ – 2)/(2*πλ*^2-*γ*^).

The kernel Eq. (9) has the same basic properties as the modified power-law kernel in Eq. (5) (Box 1; fat-tailed, power-law) but this specific form was used by Mikaberidze et al. (2016) to improve computational efficiency; we used the same function here to be able to compare the results. Similarly to Eq. (5), γ is the shape parameter, λ is the scale parameter (set to 0.762 m).

Similarly to the septoria tritici blotch case above, we first used the point source approximation, assuming both the source and the destinations to be points and interpreting the kernel Eq. (9) as a function of one spatial coordinate (i.e., set *r* = *x, k* =1). Using Eq. (9), we computed the disease severity after dispersal at a distance *x* from the source

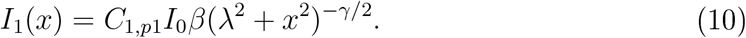

Next, we used the spatially explicit approach. We computed the disease severity (averaged over the length of a measurement line) at a distance *x_d_* from the middle of the source by substituting the kernel Eq. (9) into Eq. (1) and specifying the integrals according to experimental design

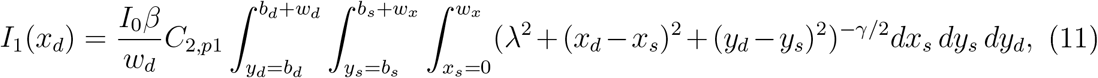

where *w_x_* = *w_y_* = 1.53 m is the side length of the square source, *b_s_* = 2.285 m is the gap between the inoculation area and the edge of the plot, *w_d_* = 3m is the length of each measurement line (along the *y*-axis), *b_d_* = 1.525 m is the width of the border excluded from measurements at each end (see Fig. 2B).

Following Sackett and Mundt (2005), and Mikaberidze et al. (2016), we performed a natural logarithmic transformation of observed disease gradients to avoid a disproportionate emphasis on the few large values at the beginning of the gradient, and excluded zeros from the log-transformed data. Accordingly, we log-transformed both the one-dimensional model Eq. (10) and the two-dimensional model Eq. (11). We then fitted both functions to log-transformed disease gradients to estimate the shape parameter *γ*

#### Potato late blight

We analyzed a subset of data on dispersal of *P. infestans* (Gregory, 1968, Table III, unsprayed experiment). *P. infestans* zoospores were inoculated across a strip in the middle of each experimental plot (orange area in Fig. 2C). Plots were 8.5m-wide and over 60 m-long, rows of potato plants were oriented along the plot length (along the *x*-axis). Disease severity was measured as numbers of late blight lesions counted visually on all leaves belonging to two adjacent potato stems chosen in every row at several distances from the source: from 1 m up to 30 m (Fig. 2C).

When describing the experiment, Gregory (1968) did not report the widths of the inoculation areas and the measurement lines (i.e., their extent along the *x*-axis), but only stated that they spanned five plants and two plants along a row, respectively. The paper also did not specify the location of the origin from which the distances between the source and the measurement locations were determined. We inferred these details based on reasonable assumptions. First, we assumed the typical distance between plants along the row to be 0.38 m (corresponding to the typical planting density of potatoes). Based on that, we computed the width of the inoculation area (i.e., the extent of five plants along a row) as 1.9 m and the width of a measurement line (i.e., the extent of two plants along a row) as 0.76 m. Second, we assumed that the reported distances were measured from the edge of the inoculation area that is closer to the gradient under consideration (Fig. 2C), as measuring them from the center of the inoculation area would lead to an overlap between the first measurement line and the inoculation area.

The power-law function

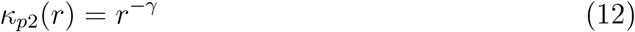

used to describe disease gradients by Gregory (1968) diverges at *r* = 0 and is not a probability density function, and hence this function is not a kernel. Nevertheless, we used it in the analysis as if it were a kernel so that we can compare the estimates of the shape parameter *γ* with the estimates obtained by Gregory (1968).

As we did earlier, we first used the point source approximation, assuming both the source and the destinations to be points. Under this approximation, we used Eq. (12) to compute the disease severity after dispersal at a distance *r* = *x* from the edge of the source

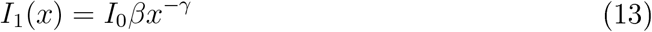

Next, we used the spatially explicit approach and computed the disease severity (averaged over the length of a measurement line) at a distance *x_d_* from the source by substituting Eq. (12) into Eq. (1) and specifying the integrals according to the experimental design

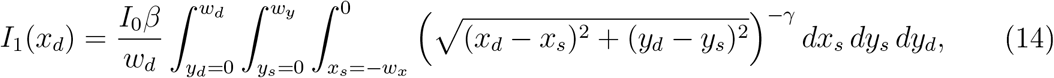

where *w_x_* = 1.9 m and *w_y_* = 8.5 m are the dimensions of the inoculation area. Both the inoculation area and the measurement lines spanned the entire width of the plot: from 0 to *w_d_* (hence, here *w_y_* = *w_d_*}. Since the function in Eq. (12) cannot be normalized, there is no normalization factor in Eq. (13) and Eq. (14). For this reason, the prefactor *I*_0_*β* in Eq. (13) and Eq. (14) has no biological relevance.

We performed the natural logarithmic transformation of observed disease gradients and excluded zeros from transformed data. Accordingly, we log-transformed both the one-dimensional Eq. (13) and two-dimensional Eq. (14). Then, we fitted the two functions to observed disease gradients to estimate the shape parameter γ.

#### Data analysis

The fitting was implemented in Python 3.7 using packages numpy (v. 1.17.3, Harris et al., 2020), scipy (v. 1.3.1, Virtanen et al., 2020) and lmfit (v. 1.0.1, Newville et al., 2014). Fitting was performed as a least squares optimization using function ’Model.fit’ of package lmfit, which in turn uses either Levenberg-Marquardt algorithm or Trust Region Reflective algorithm in the function scipy.optimize.least_squares. The same fitting method was used in all analyses of the experimental data in this section and in numerical simulations in the following section. In the analysis of experimental data, we also used the brute force optimization over a grid to achieve reasonable starting values for the least squares optimization.

We used the estimates of kernel parameters obtained from the fitting procedure for septoria tritici blotch and stripe rust to quantify the characteristic scales of dispersal by computing medians (*r*_50_) and 90th percentiles (*r*_90_) of dispersal distance kernels (Nathan et al., 2012). We computed the two percentiles numerically by solving the equation

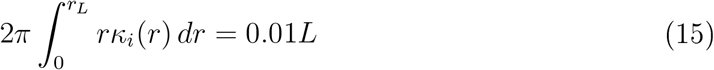

with respect to *r_L_* at *L* = 50,90. Here, *κ_i_*(*r*) is the dispersal kernel function, where *i* = *e,p*1; *e* stands for the exponential kernel [Eq. (3) at *k* = 2] and *p*1 stands for the modified power-law kernel [Eq. (9) at *k* = 2], The two percentiles, *r*_50_ and *r*_90_, correspond to the distances up to which 50% and 90% of the dispersed population has advanced.

### Results of the analysis

We fitted the measured dispersal gradients in the three case studies to model functions using the spatially explicit approach (2D-estimation) and the point source approximation (1D-estimation; see Fig. 3, left column). In all three examples, 2D-estimation resulted in steeper dispersal kernels compared to 1D-estimation (cf. solid blue and dashed orange curves in the right column of Fig. 3). This is because the estimates of kernel parameters differed between 1D- and 2D-estimation approaches (Table 1). We observed a moderate reduction of about 12% in the *α*-estimate for septoria tritici blotch, a moderate increase of about 10% in the γ-estimate for stripe rust, and a more substantial increase exceeding 30% in the γ-estimate for potato late blight (Table 1). The goodness of fits, measured as sums of squared residuals (SSR in Fig. 3), did not differ much between the two methods, but slightly favored the 1D fit.

**Figure 3:**
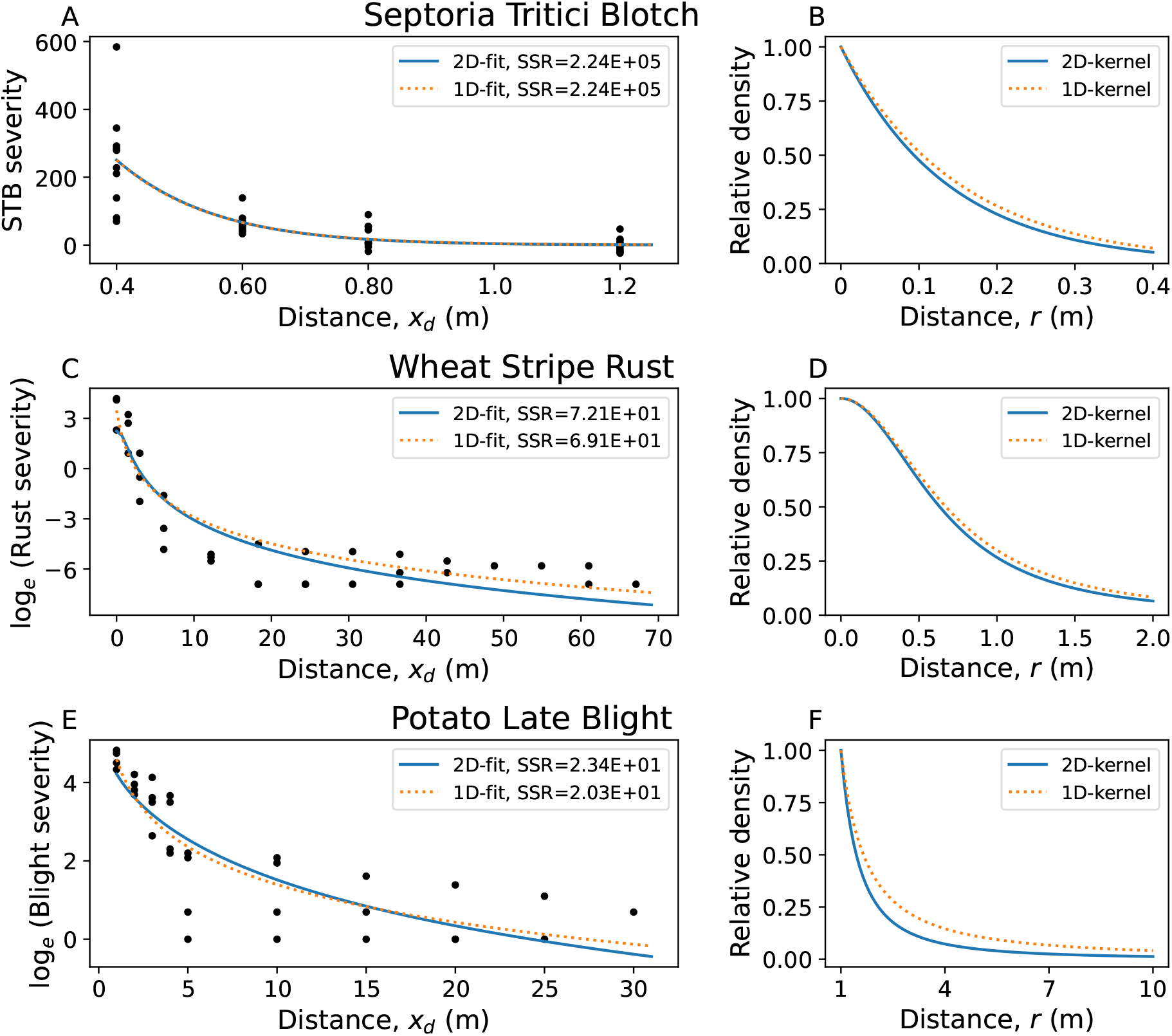
Estimation of dispersal kernels from empirical datasets. Left column: observed disease gradients (black circles) and the best-fitting models based on the spatially explicit approach (“2D-fit”, solid blue curves) and based on the point source approximation (“1D-fit”, dotted orange curves). The legends display the sums of squared residuals (SSR). Right column: the estimated dispersal kernels with best-fitting parameters according to the spatially explicit approach (“2D-kernel”, solid blue curves) and the point source approximation (“1D-kernel”, dotted orange curves). Kernels are normalized to start from one at *r* = 0 m for septoria tritici blotch and stripe rust but at *r* = 1 m for late blight, as the chosen power-law function is not defined at zero.

**Table 1:**
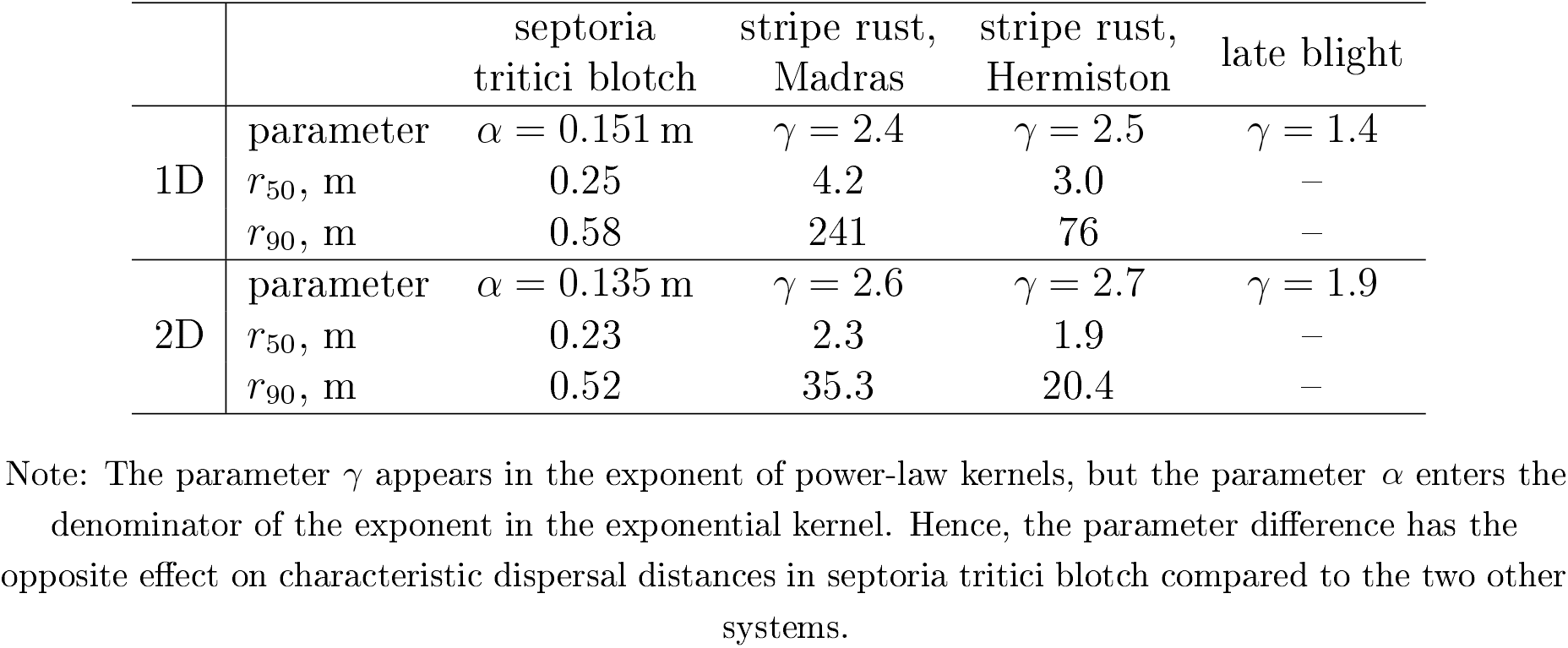
Comparison of kernel parameter estimates and associated percentiles of dispersal distance kernels between one- and two-dimensional models (1D and 2D, respectively).

For septoria tritici blotch and stripe rust, we also investigated how the differences in kernel parameter estimates affect the characteristic spatial scales of dispersal, quantified by the 50th and 90th percentiles of dispersal distance kernels, *r*_50_ and *r*_90_. For septoria tritici blotch, a moderate reduction in the estimate of the kernel parameter *α* according to 2D-estimation compared to 1D-estimation translated into a similarly moderate decrease in *r*_50_ and *r*_90_ (2nd column in Table 1). In contrast, for stripe rust, a modest increase of about 10% in the estimate of the kernel parameter γ translated into a dramatic decrease in the spatial scales of dispersal (3rd and 4th columns in Table 1). For example, for Madras dataset, 2D-estimation resulted in a nearly two-fold reduction of *r*_50_ and a massive, almost seven-fold reduction of *r_90_* (3rd column in Table 1). We expect a comparably strong reduction in estimated characteristic scales of dispersal for potato late blight, but could not conduct this analysis with the power-law function defined by Eq. (12), because it cannot be normalized. Thus, according to 2D-estimation, spores of all three organisms move on average over shorter distances compared to 1D-estimation. This reduction is more prominent in stripe rust compared to septoria tritici blotch.

## Simulations

In the case studies above, the spatially explicit approach (2D-estimation) resulted in substantially different estimates of dispersal kernels compared to the point source approximation (1D-estimation). Are these estimates more accurate, i.e., closer to the true values? This is plausible, because the 2D-estimation describes dispersal from spatially extended sources more realistically. However, we cannot answer this question definitively based on the analysis of experimental data alone, because we do not know the true values of dispersal kernel parameters. Here, we addressed this question via numerical simulations. We first simulated the dispersal process according to exponential, Gaussian and power-law kernels with pre-defined parameters. Then, we used both methods to estimate the kernel parameters and compared the two methods in terms of their estimation accuracy.

We started by conducting idealized simulations: the 2D-estimation provided perfectly accurate estimates, while 1D-estimation exhibited substantial errors. We analyzed how these errors depend on the parameters of kernel functions and source sizes. Further, we considered how different parts of the extended source area ’distort’ the parameter estimates when assuming a point source. Finally, we examined a more realistic scenario that incorporated extended measurement areas (instead of measurement points) and a limited amount of sampling within the areas.

### Methodology of simulations

#### Which parameters did we estimate?

The exponential and Gaussian kernel functions have a single parameter: the scale parameter *α* that characterizes the spatial scale of dispersal (see Box 1) and this was the parameter we estimated. In contrast, the power-law kernel has two parameters: the scale parameter A and the shape parameter γ both of which influence the spatial scale of dispersal. To make the estimation comparable between the three different kernels, we set A to its true value and estimated only γ. (We estimated both parameters in Appendix A). Thus, we estimated a single parameter in each of the three dispersal kernel functions, which we call the “kernel parameter”.

#### Idealized simulations

We simulated dispersal from a spatially extended source and sampled the resulting dispersal gradients in an idealized manner: we assumed that sampling locations were points without any spatial extent and that the measured values accurately reflected the true values.

First, we fixed the source size (1 m × 1 m) and conducted simulations at different mean dispersal distances from 1 m to 50m (in 1 m steps). (Expressions for mean dispersal distances for each of the three kernels are given in Box 1). Next, we simulated dispersal with a fixed mean dispersal distance (20 m), but varied the size of the square source from 1 m × 1 m to 30m × 30m (in 1m steps).

We derived the number of dispersed individuals from Eq. (1) by considering the destination as a point and using a square source area:

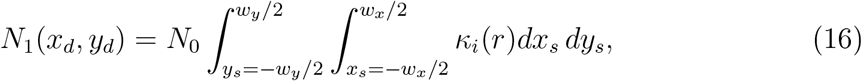

where the origin is set to the center of the square source, 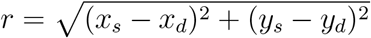 is the distance between the source point (*x_s_, y_s_*) and the destination point (*x_d_, y_d_*), *w_y_* = *w_x_* is the side length of the square source, and *κ_i_*(*r*) is the dispersal kernel function, where *i* = *e,g,p* stands for exponential, Gaussian and power-law kernels, respectively (we used Eq. (3)-(5) in Box 1, setting *k* = 2, and setting λ = 0.5m for the power-law kernel). We assumed in Eq. (16) that the population of *N*_0_ dispersing individuals was uniformly distributed across the spatial extent of the source.

We generated the dispersal gradients by evaluating the number of dispersed individuals *N*_1_(*x_d_, y_d_*) according to Eq. (16) at *y_d_* = 0 and a number of different values of *x_d_*, starting from 10 cm away from the edge of the source in steps of 10 cm until reaching the point where the number of dispersed individuals was less than 1% of that of the first measurement point (dotted lines in Fig. 4A and 4B).

**Figure 4:**
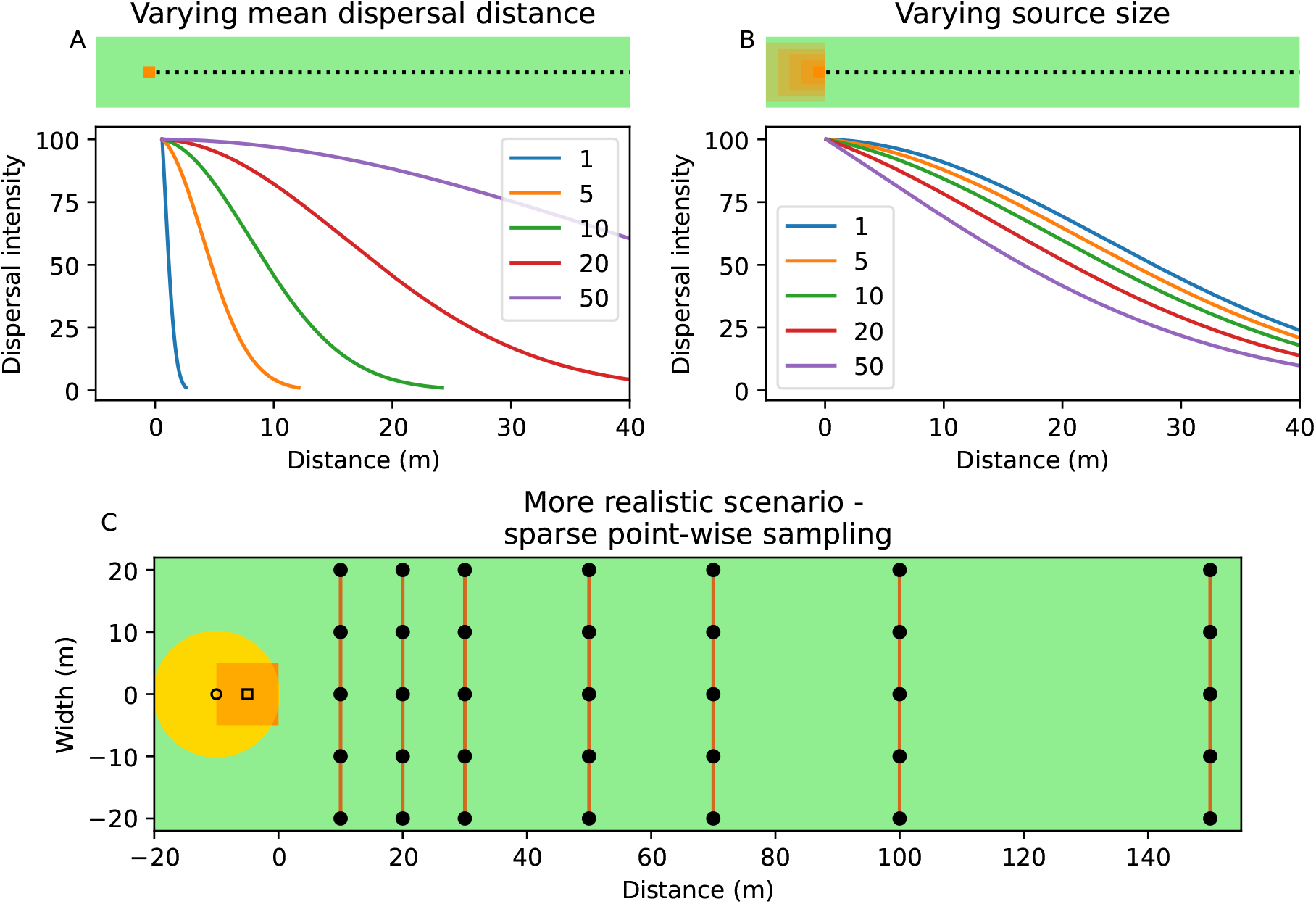
Design of simulations. A) and B) plot designs and example gradients in the idealized simulations with the Gaussian kernel Eq. (4). Measurements were taken along the dotted line every 10 cm. A) Mean dispersal distances were varied from 1 m to 50 m, while the size of the square source was fixed to 1 m × 1 m. B) Source side lengths were varied from 1 m to 50 m, while the mean dispersal distance was fixed to 20 m. C) A more realistic scenario considered two source shapes (orange circle and square) and sparse, point-wise sampling, where black circles show measurement locations (see text). Black open circle and square represent virtual point sources at the centers of the actual source areas.

The kernel parameter was then estimated from simulated data (i) using a two-dimensional dispersal kernel and incorporating the spatial extent of the source according to Eq. (16) with the kernel functions given by Eq. (3)-(5), setting *k* = 2; and (ii) using the corresponding one-dimensional dispersal kernels (Eq. (3)-(5) in Box 1, setting *k* =1), where we set *r* = *x_d_* in the expressions for the kernel functions (i.e., considering a virtual point source located at the center of the actual source area). In Appendix A, we used a similar analysis to explore the case when the virtual point source is located at the edge of the source area, rather than its center (as it is sometimes assumed in dispersal experiments, for example, in the potato late blight case study above).

Next, we investigated how individual points within the source area influence the accuracy of the point-source approximation. For this purpose, we considered a square 1 m × 1 m source and a dispersing population with 2m mean dispersal distance. We simulated dispersal from each individual point within the source area (over a 2 cm × 2 cm square grid) for each of the three kernel functions (setting λ = 0.5 m for the power-law kernel). In this way, we generated the dispersal gradients produced by individual points across the area of the source according to Eq. (16). Then, we estimated the kernel parameters (as described above) and the associated mean dispersal distances assuming that the source was a point located at the center of the source area. We then recorded whether the estimates of kernel parameters were higher or lower than their true values (Fig. 6).

#### More realistic simulations

We considered a scenario that incorporated key features of real dispersal experiments: not only a spatially-extended source, but also a spatially-extended destination and a limited sampling density. As we did earlier, we simulated dispersal governed by the three different kernel functions. The kernels and their parameters are given in Fig. 1: exponential, Gaussian and power-law kernels with mean dispersal distance 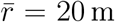 (λ = 20m for power-law kernel). In addition to the square source area (10m × 10m), here we also considered a circular source area (20 m diameter). Dispersal was simulated in a deterministic manner similarly to the previous subsection: we evaluated the number of dispersed individuals at a number of different values of *x_d_*, corresponding to distances of 10 m, 20 m, 30 m, 50 m, 70 m, 100 m and 150 m from the edge of the source that is adjacent to measurement lines (Fig. 4C). In contrast to idealized simulations above, we also incorporated the spatial extent of destination areas: we considered the destinations as thin lines along the *y*-axis (“measurement lines”, brown vertical lines in Fig. 4C). To achieve this, we evaluated *N*_1_(*x_d_, y_d_*) according to Eq. (16) for each value of *x_d_* (i.e., for each measurement line) at five different values of *y_d_* evenly spaced across the length *w_d_* = 40m of the measurement line (filled black circles in Fig. 4C). We investigated the role of increasing the number of sampling points from five to 10, 40, and 160, and the role of shortening the measurement lines from 40 m to 20 m, 5 m, and 1 m in Appendix B.

In a typical dispersal experiment (such as the ones considered above in “Analysis of empirical data” section), a number of measurements are conducted across measurement lines and their outcomes are recorded together with the value of *x_d_*, but it is not practically feasible to also record the values of *y_d_.* For this reason, an average is taken over all measurements conducted within each measurement line and these average values of *N*_1_ versus *x_d_* represent the measured dispersal gradients. We mimicked this measurement process here by computing an average over all *N*_1_-values within each of the measurement lines.

We fitted the resulting dispersal gradients to models to estimate kernel parameters in the following way. First, the spatially explicit fitting was performed, whereby the function

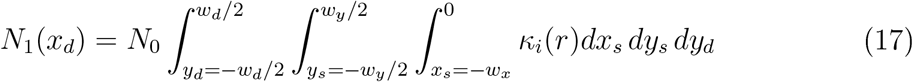

that describes the number of dispersed individuals *N*_1_(*x_d_*) versus *x_d_* was fitted to simulated dispersal gradients. The meaning of variables and parameters in Eq. (17) is the same as in Eq. (16). Here, we also conducted integration over the length of the measurement line *w_d_* (the outer integral in Eq. (17)). In doing this, we assumed that the number of measurements taken within each measurement line was sufficiently high.

Second, the fitting under the point-source approximation was performed using one-dimensional kernels (Eqs. (3)-(5) in Box 1, setting *k* = 1), where we set *r* = *x_d_* + 5m for the square source and *r* = *x_d_* + 10 m for the circle source in the expressions for the kernel functions, thereby considering a virtual point source located at the center of the actual source area.

### Results of simulations

#### How does the accuracy of the point-source approximation depend on the kernel parameter and the source size?

In the idealized simulations, where we incorporated the spatial extent of the source (2D-estimation), the kernel parameter estimates were error-free in all cases (up to very small errors inherent in numerical computation). In contrast, the estimation under the assumption of a point source (1D-estimation) resulted in substantial errors. We investigated how these errors change when we vary the true values of mean dispersal distances (Fig. 5A) and when we vary the source size (Fig. 5B). We also estimated mean dispersal distances and investigated how errors of these estimates depend on the true values of mean dispersal distances (Fig. A1C in Appendix A) and on the source size (Fig. A1D in Appendix A).

**Figure 5:**
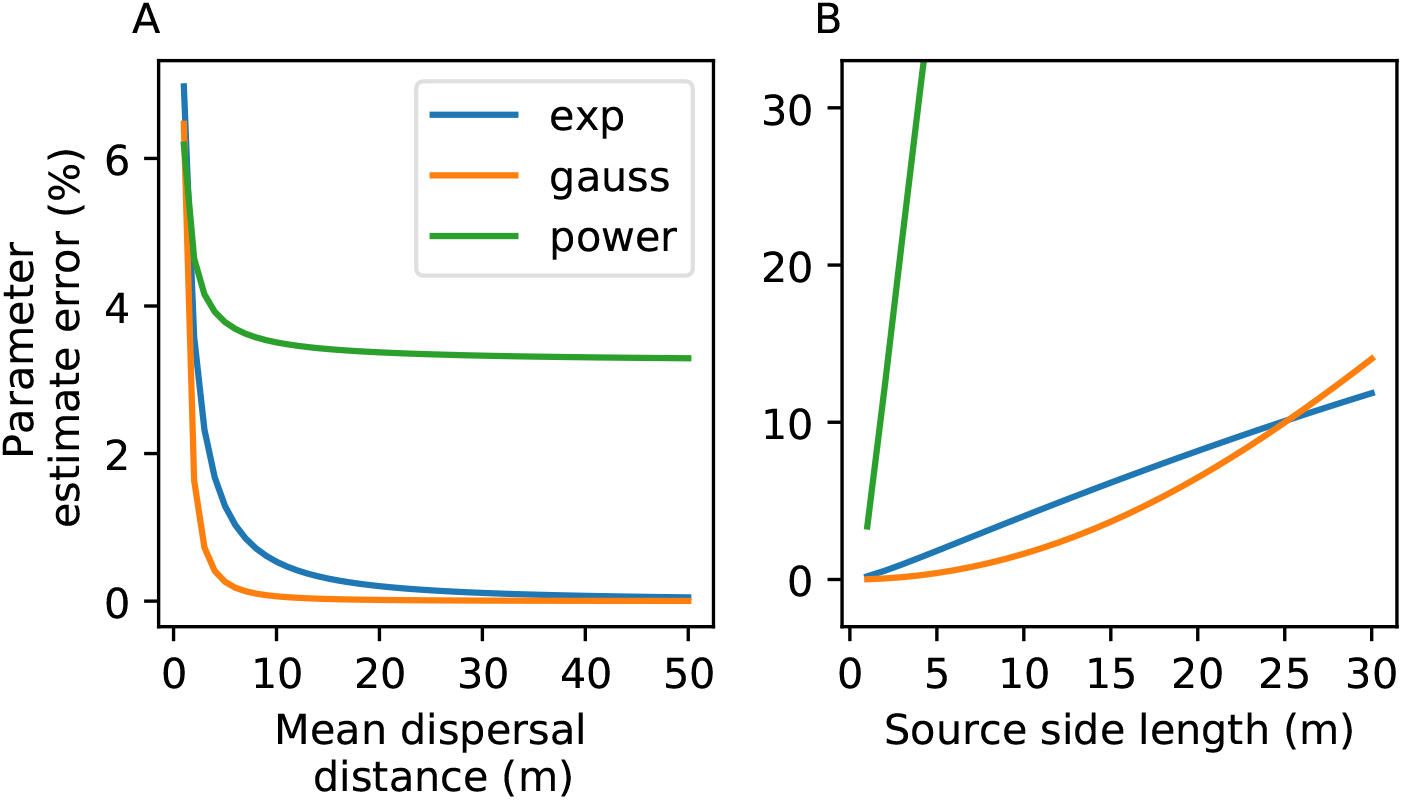
Effect of the mean dispersal distance (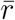, left panel) and source size (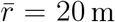, right panel) on the accuracy of the point source estimate (assuming a point source at the center of the actual square source). The error in the kernel parameter estimates is shown as a function of true mean dispersal distance and source size. Simulation design corresponds to Fig. 4A,B. Accuracy of estimation improves when considering organisms with longer mean dispersal distances and when using smaller sources. Parameters: *βI*_0_ = 10000 (arbitrary), λ = 0.5m, 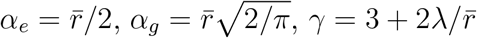.

The estimates of the kernel parameter become more accurate at longer mean dispersal distances for each of the three kernel functions (Fig. 5A). Similarly, the estimates become more accurate when using smaller sources (Fig. 5B).

Estimates of the power-law kernel parameter behave differently compared to the two other kernel functions. First, for both exponential and Gaussian kernels, the errors approach zero when the mean dispersal distance is increased, but for the power-law kernel, the errors appear to approach a substantially positive value (cf. green curve with blue and orange curves in Fig. 5A). Second, estimation errors increase with the size of the source much faster in the case of the power-law kernel compared to exponential or Gaussian kernels (cf. green curve with blue and orange curves in Fig. 5B).

In all three kernel functions, the point-source approximation leads to an overestimation of their respective kernel parameters (all errors have a positive sign in Fig. 5). However, the mean dispersal distances are overestimated for exponential and Gaussian kernels but underestimated for the power-law kernel under the point-source approximation. This difference arises because the mean dispersal distance is proportional to the kernel parameter *α* in the case of exponential and Gaussian kernels, but inversely proportional to the kernel parameter γ for the power-law kernel. (See Fig. A1C,D in Appendix A for the investigation of errors in estimation of mean dispersal distances.)

To conclude, when we estimate the kernel parameter under the point source approximation, the estimation accuracy increases for organisms with longer mean dispersal distances and when using smaller source sizes. Thus, in this idealized scenario, the estimation of kernel parameters under the point-source approximation could reach any desired level of accuracy by using a sufficiently small source.

#### How is the accuracy of the point-source approximation influenced by the spatial extent of the source?

To better understand the patterns we observed in simulations above, we studied in more detail how the errors in the estimation of kernel parameters arise under the point-source approximation. (A similar investigation of the errors in estimation of mean dispersal distances is presented in Appendix A). We decomposed the overall error associated with the point-source approximation into its more basic ingredients: we considered how each displacement of the point source from its location at the center of a square source to a different location within the source area contributes to the error in the estimated kernel parameter.

When dispersal is governed by an exponential kernel, we observe a simple pattern. Points within the source area along the continuation of the line on which measurements are conducted (the horizontal line drawn through the center of the source area in Fig. 6A) yield accurate estimates, whereas any other points within the source area result in an overestimation of the kernel parameter. This follows from the memorylessness property of exponential kernels (Box 2): dispersal gradients produced by the points along this line all have the same shape and this shape corresponds exactly to the shape of the gradient produced by the point source at the center of the source. Any displacement of the source point from this line leads to a modification of the gradient shape (more specifically, to flattening of the gradient) and hence an overestimation of the kernel parameter.

**Figure 6:**
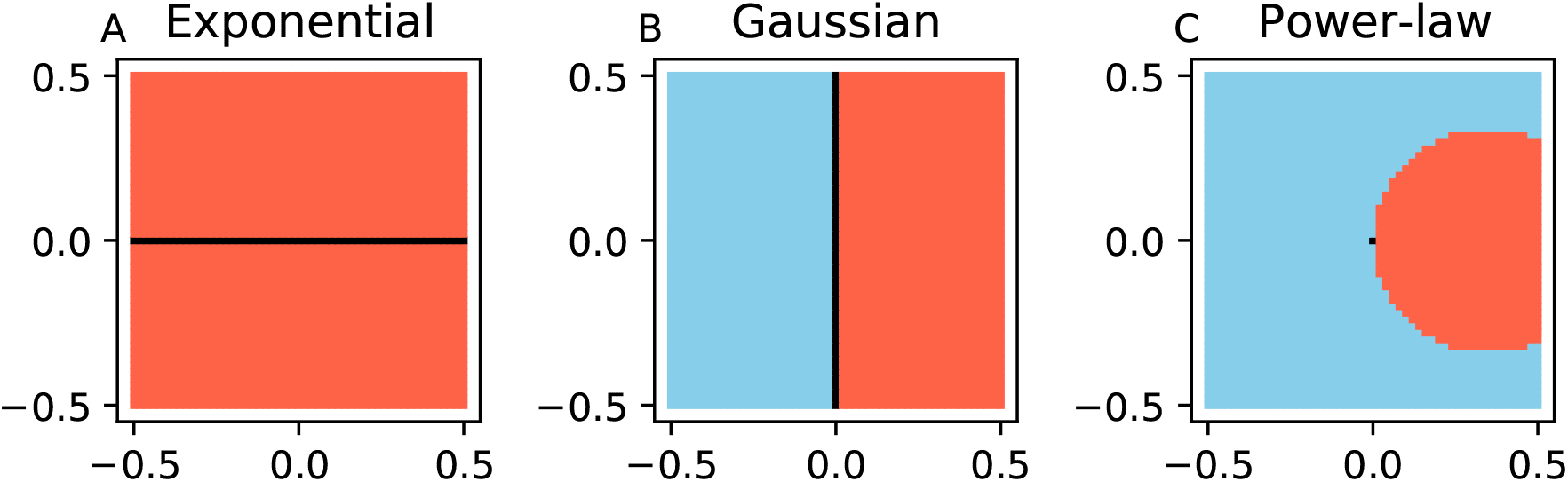
Contributions of individual points within the 1 m × 1 m source area to the errors in the kernel parameter estimates. Here, we considered a virtual pointsource at the center of the actual square source area. For each point within the source area (across a 2 cm × 2 cm grid) the color shows, whether the point contributes to an underestimation (blue) or an overestimation (red) of the kernel parameter, or represents an accurate estimate (black). Measurements are conducted on the right side of the source area as in Fig. 4A. Parameters: *βI*_0_ = 1 (arbitrary), 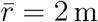, λ = 0.5 m, 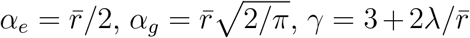.

Gaussian kernel exhibits a similarly simple pattern that stems from its separability property (Box 2). In this case, the line drawn through the center of the source perpendicularly to the line of measurement locations has a special significance (Fig. 6B). Each point source on this line produces a gradient of the same shape as the point source at the center of the source area, leading to an accurate estimation of the kernel parameter. Any other points within the source area produce either steeper gradients (left half of the source area) or flatter gradients (right half of the source area). This leads to an underestimation (blue area in Fig. 6B) or an overestimation (red area in Fig. 6B) of the kernel parameter.

The power-law kernel is neither memoryless nor separable and hence we observe a more complex pattern: points within a circular area adjacent to the measurement locations lead to an overestimation of the kernel parameter, while all other points within the source area result in its overestimation (Fig. 6C).

Based on these patterns in Fig. 6 we can now understand why the point-source approximation leads to an overestimation of kernel parameters for all three kernel functions in Fig. 5 above. The overall gradient produced by the square source is a weighted sum over individual gradients produced by each of the point sources comprising the spatially-extended source (according to Eq. (16)). Under the point-source approximation, we estimate the kernel parameter by fitting the one-dimensional kernel function to this overall gradient. Hence, the error of this estimation can be understood on the basis of the errors resulting from displacing the point source from the center to a different location within the source area.

For the exponential kernel, every such displacement produces a dispersal gradient, which when fitted by a one-dimensional kernel results in an overestimation of the kernel parameter: the whole area of the source is red in Fig. 6A, except for the black horizontal line. Therefore, fitting of the one-dimensional kernel function to the overall dispersal gradient will also lead to an overestimation of the kernel parameter. In the case of the Gaussian kernel, one half of the displacements of the point source from the center lead to an overestimation, while the other half leads to an underestimation of the kernel parameter (Fig. 6B). But when we compute the overall gradient as a weighted sum over individual gradients produced by each of the points sources, and estimate the kernel parameter under the point-source approximation, the kernel parameter is overestimated. This is because the point sources that lead to an overestimation (the right half of Fig. 6B) contribute with higher weights, since they are closer to the measurement locations. In the case of the power-law kernel, a larger part of the source area corresponds to displacements of the point source leading to an underestimation of the kernel parameter (the blue region outside of the red circular area in Fig. 6C). Yet again, the displacements of the point source that lead to an overestimation of the kernel parameter (those within the red circular area in Fig. 6C) make disproportionately high contributions since they are closer to the sampling locations. For this reason, the fitting of the one-dimensional kernel function to the overall gradient generated by the whole source area leads to an overestimation of the kernel parameter.

Thus, by analyzing the effects individual displacements of the point source from the center of the source area, we gained a deeper understanding of how the point source approximation produces errors in estimates of dispersal kernel parameters.

#### Estimation of dispersal kernel parameters in a more realistic scenario

So far we considered idealized simulations, where we found that errors in estimation of kernel parameters stem entirely from the point-source approximation. Here, we investigate a more realistic scenario that accounts for the fact that sampling locations are distributed over a finite area and sampling is conducted a limited number of times.

The spatially explicit estimation resulted in more accurate estimates of the kernel parameters in all cases (two different sources and three different kernels), but not entirely accurate (Table 2). We found that the reason for this remaining error was the finite number of sampling points within the extended destinations (i.e., along the measurement lines). When we increased the number of sampling points, the spatially explicit estimates approached their true value, while the estimates acquired under the pointsource approximation retained a substantial error (Appendix B).

**Table 2:** Estimation of kernel parameters from simulated dispersal data. The spatially explicit approach (“2D”) provides more accurate estimates.

| Source | Kernel | Parameter | 1D | 2D | True |
| --- | --- | --- | --- | --- | --- |
| Square | Exponential | $\alpha$ | 11.34 | 10.26 | 10.0 |
| | Gaussian | $\alpha$ | 16.25 | 16.00 | 16.0 |
| | Power-law | $\gamma$ | 4.27 | 4.83 | 5.0 |
| Circle | Exponential | $\alpha$ | 11.11 | 10.21 | 10.0 |
| | Gaussian | $\alpha$ | 16.77 | 16.00 | 16.0 |
| | Power-law | $\gamma$ | 4.54 | 4.85 | 5.0 |

Note: “2D” and “1D” stand for two- and one-dimensional models, respectively; $\alpha$ is measured in m, $\gamma$ is unitless. The scale parameter of the power-law kernel is set to $\lambda = 20$ m.

Interestingly, for exponential or Gaussian kernels, the point source approximation leads to overestimation of kernel parameters both here (Table 2, column “1D”) and in the idealized simulations in Fig. 5. In contrast, for the power-law kernel, the kernel parameter is underestimated here (Table 2, column “1D”), but overestimated in Fig. 5. Why?

To find out, we conducted additional simulations, where we varied the length of measurement lines (brown vertical lines in Fig. 4C). We found that the error in the estimate of the power-law kernel parameter changes its sign when the measurement lines become sufficiently long (Appendix B; Table B3).

To conclude, the spatially explicit estimation remains substantially more accurate than the point source approximation in more realistic simulations. For the power-law kernel, the estimation error caused by the point source approximation depends in a non-trivial manner not only on the spatial scale of dispersal and the source size (Fig. 5), but also on the extent of measurement lines (Appendix B; Table B3). This makes it difficult to devise experimental designs that lead to a sufficiently accurate estimation when assuming a point source. Thus, we presented compelling evidence in favor of adopting the spatially explicit approach to estimation of dispersal kernel parameters.

## Discussion

We devised a theoretical framework that allows to extract knowledge on dispersal kernels from empirical dispersal gradients by incorporating the spatial extent of dispersal sources. We re-analyzed existing dispersal gradients for three major plant pathogens and found that this spatially explicit approach provides considerably different estimates of dispersal kernels compared to the conventional point source approximation. Further, we demonstrated using numerical simulations that the spatially explicit approach yields more accurate estimates in a wide range of biologically plausible scenarios. Combining these two lines of evidence, we conclude that the three organisms disperse on average over substantially shorter distances compared to estimates from conventional modeling. Using this method, a significant proportion of published dispersal gradients (e.g., Werth et al., 2006; Skarpaas and Shea, 2007; Loebach and Anderson, 2018; Emsweller et al., 2018; D’Aloia et al., 2015; Devaux et al., 2006) can be re-analyzed to improve our knowledge about spatial scales of dispersal. Thus, our results can boost progress in empirical characterization of dispersal across different taxa.

The theory we formulated here is not new, and similar spatially explicit approaches were used in modeling studies to investigate dispersal in plants (Clark et al., 1999; Shaw et al., 2006). However, such approaches are not adopted in the literature on empirical characterization of dispersal (e.g., not used in Werth et al., 2006; Skarpaas and Shea, 2007; Loebach and Anderson, 2018; Emsweller et al., 2018; D’Aloia et al., 2015; Devaux et al., 2006). Also, Bullock et al. (2017) excluded dispersal gradients produced by line and area sources from their analysis, because they could not be compare them to dispersal gradients from point sources. Dispersal kernels estimated using the spatially explicit approach presented here enable such comparisons, because the estimates are independent of specific experimental design. The novelty of our study lies in the comprehensive demonstration of how to use this theory to extract more knowledge from existing datasets on dispersal, enabling more meaningful comparisons between different experiments.

We assumed isotropic dispersal in data analysis and simulations. However, anisotropic dispersal is common in nature (Soubeyrand et al., 2007) and the model can be extended to incorporate it. In this extended model, the probability of dispersal from a source point to a destination point will depend not only on the distance between the points, as in our case, but also on the direction from the source to the destination. Parameters of anisotropic dispersal kernels can then be estimated from measurements of dispersal gradients with the spatially explicit consideration of the source. Furthermore, we focused on two-dimensional models of dispersal, which are simplifications of the three-2D versusdimensional dispersal process (?). If the third dimension is of importance in the system, such as for aquatic environments, tree canopies (Cousens and Rawlinson, 2001) or crop canopies (Vidal et al., 2018), a three-dimensional model may be needed to appropriately describe dispersal. Empirical data we analyzed here was collected in designed experiments and characterized populations of passively dispersing organisms. However, a similar approach to data analysis can also be used in observational studies in nature and to study actively dispersing organisms, provided that two conditions are fulfilled. First, it should be possible to clearly identify and measure the source area; and second, the dispersal process should be adequately described by means of dispersal kernels.

The methodology and the outcomes presented here can inform design of future dispersal experiments. The spatially explicit approach to estimating dispersal kernel parameters allows to use substantial source sizes, comparable with the characteristic spatial scale of dispersal, but maintain higher estimation accuracy compared to the point source approximation (2D versus 1D in Table 2). Hence, larger sources can be used deliberately to boost the output from the source and consequently the availability of samples to record, which is often a limiting factor.

Based on the outcomes of our idealized simulations, it is tempting to propose simple rules of thumb about when the point source approximation provides reasonably accurate estimates of dispersal kernel parameters. This appears to be the case, for example, when the sources are sufficiently small and the spatial scale of dispersal is sufficiently large (Fig. 5). However, in these idealized simulations, we neglected the spatial extent of measurement areas and limitations in the amount of sampling within these areas. When we considered these features in more realistic simulations, the outcomes revealed a nontrivial influence of several factors (such as the functional form of the kernel, the spatial configuration of the source and the measurement locations as well as sample sizes) on the estimation accuracy. As a result, we are not able to provide simple rules of thumb regarding the validity of the point source approximation.

Instead, our results suggest the following best practices for the design of future dispersal experiments. First, a proposed experiment should be simulated numerically over a range of plausible parameter values. This will help to decide whether the point source approximation is valid or the spatially explicit modeling should be used in the analysis. Second, aspects of experimental design, such as the spatial configurations of the source and the measurement areas and the sample sizes, can be optimized by doing further simulations in order to minimize the costs of the experiment while maximizing the estimation accuracy.

## Acknowledgements

PK and AM gratefully acknowledge financial support from the Swiss National Science Foundation through the Ambizione grant PZ00P3_161453.

## Data and Code Accessibility Statement

The source code together with data is provided in a data repository (Data dryad, TBA).

## Appendix A: Further analysis of errors associated with the point-source approximation

### Accuracy of estimation of mean dispersal distances

In the “Simulations” section of the main text, we explored how the accuracy of estimation of kernel parameters under the point-source approximation. Here, based on the estimates of kernel parameters, we also estimated the mean dispersal distances and investigated how errors of these estimates depend on the true values of mean dispersal distance (Fig. A1C) and on the source size (Fig. A1D).

When dispersal is governed by exponential or Gaussian kernels, increasing the mean dispersal distance leads to a more accurate estimation of the kernel parameter. This translates into more accurate estimation of the mean dispersal distance (cf. blue curves and orange curves between Fig. A1A and A1C). In contrast, when dispersal is governed by the power-law kernel the increased accuracy of estimation of the kernel parameter does not translate into more accurate estimates of the mean dispersal distance. Instead, the error in estimates of the mean dispersal distance grows as a function of the true value of the mean dispersal distance (Fig. A1C) and as a function of the source side length (Fig. A1D). This seemingly counter-intuitive relationship follows from the expression of the mean dispersal distance for the power-law kernel (Box 1) 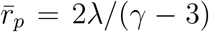. To achieve longer mean dispersal distances in Fig. A1A,C, the true value of the kernel shape parameter γ needs to become closer to the critical value of three, where the mean dispersal distance experiences a singularity. For this reason, even decreased errors in the estimates of the kernel parameter γ translate into larger errors in the estimates of the mean dispersal distance 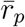.

Figure 6 in the main text showed how displacements of the point source from its location at the center of a square source to a different location within the source area contribute to the error in the estimated kernel parameter. Here, we present an analogue of Fig. 6 that considers the estimation of the mean dispersal distances in Fig. A2. Since the mean dispersal distance is proportional to the kernel parameter *α* for exponential and Gaussian kernels (Box 1), we observe the same patterns in the sign of the error when estimating the kernel parameter and the mean dispersal distance (cf. Fig. 6A,B with Fig. A2A,B). In contrast, in the case of the power-law kernel, the mean dispersal distance, 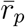, has an inverse relationship with the kernel parameter γ (Box 1). For this reason, the sign of the error in the dispersal distance estimates reverses compared to the sign of the error in the kernel parameter estimates (cf. Fig. 6C with Fig. A2C).

**Figure A1:**
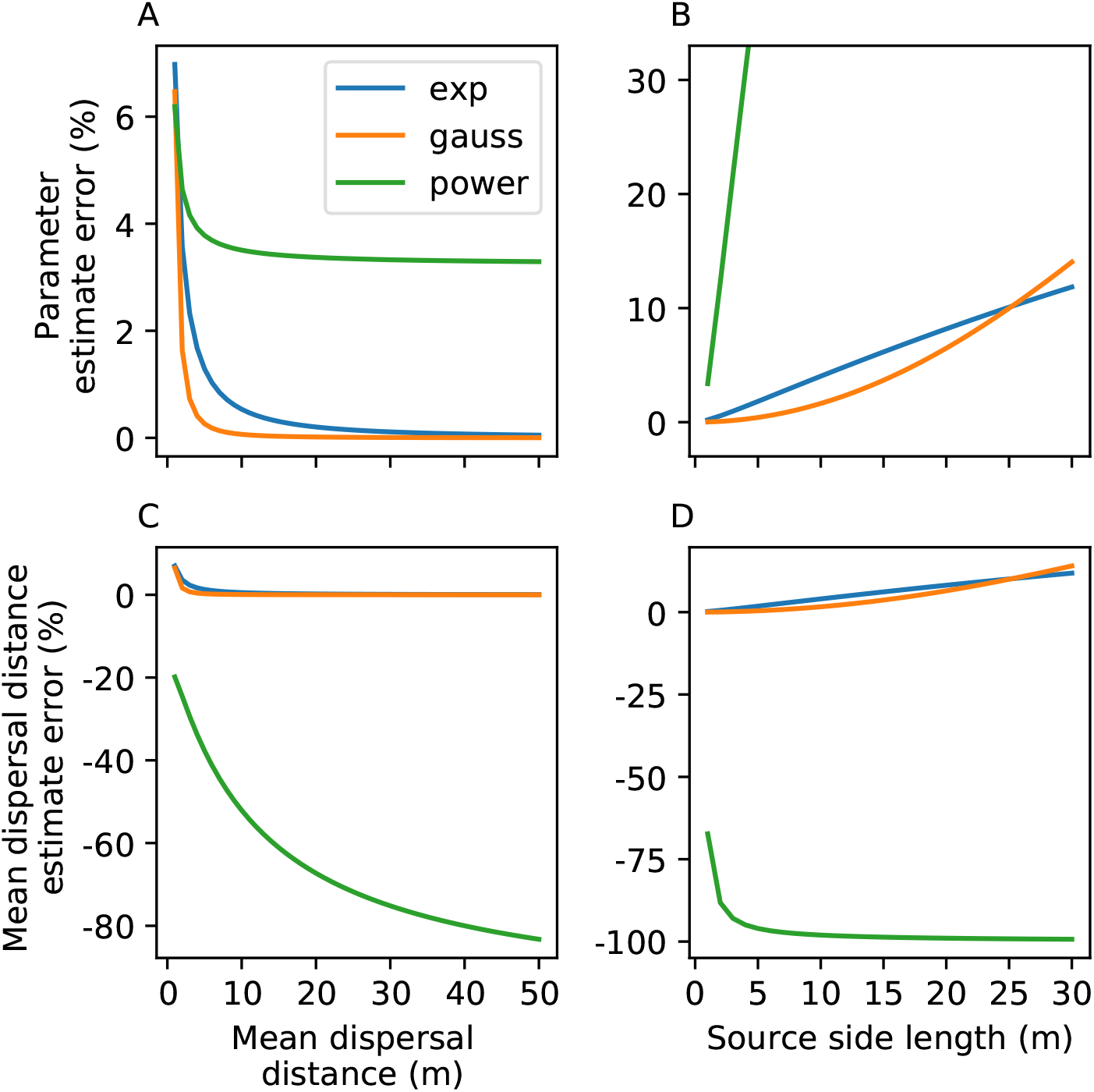
Accuracy of 1D-estimates depends on the true value of the mean dispersal distance (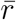, left column) and on the source size (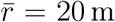, right column). Here, we considered a virtual point source located at the center of the actual square source area. Panels A and B are identical to Fig. 5 in the main text. Parameter estimates become more accurate when considering organisms with longer mean dispersal distances and when using smaller sources. Parameters: *βI*_0_ = 10000 (arbitrary), λ = 0.5 m, 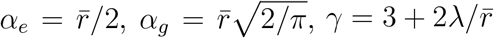.

**Figure A2:**
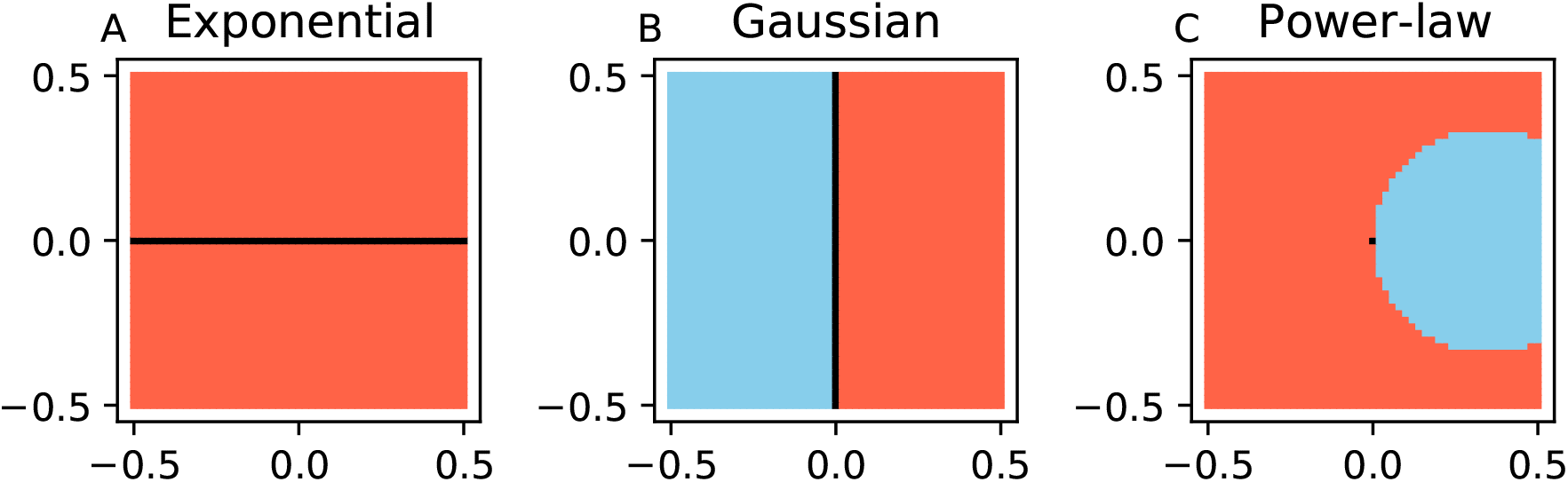
Contributions of individual points within the source area to the errors in the estimated mean dispersal distances 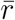. Here, we considered a virtual point source located at the center of the actual square source area. For each point within the source area (across a 2 cm × 2 cm grid) the color shows whether the point contributes to an underestimation (blue) or an overestimation (red) of the mean dispersal distance, or represents an accurate estimate (black). Measurements are conducted on the right side of the source area. Parameters: *βI*_0_ = 1 (arbitrary), 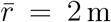, λ = 0.5m, 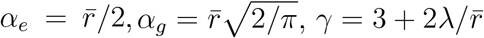.

### Virtual point source at the edge of the source area

In the main text, we investigated estimation errors associated with the point source ap-proximation considering a virtual point source located at the center of the square source area. Here, we conducted a similar analysis but considered a virtual point-source located at the edge of the source area that is closer to measurement locations. This change did not affect the overall pattern in the estimation accuracy of the kernel parameter. The estimates of the kernel parameter become more accurate when considering populations with longer mean dispersal distances and when using smaller sources (FigA3A, A3B). However, whether the mean dispersal distances are underestimated or overestimated, depends on the kernel function: they are underestimated for Gaussian kernels, but over-estimated for exponential and power-law kernels (Fig A3C, A3D). Hence, for Gaussian and power-law kernels, the error changes its sign when we move the virtual point source from the center to the edge of the source area.

We can understand this change by comparing Fig. A2 and Fig. A4. In Fig. A2 the points to the right of the middle are closer to the measured gradient and hence have a more prominent effect on the shape of the overall gradient. Therefore, the right-hand side (red) area in Fig. A2B and the blue circular area in Fig. A2C contribute to the overall gradient with larger weights and result in the over estimation of the mean dispersal distance for the Gaussian kernel but underestimation for the power-law kernel (as seen in Fig. A1C, D). However, when we move the virtual point source to the edge of the source area, the measurements of the gradient start closer to the virtual point source point and those above mentioned areas of the source no longer contribute to the gradient. Hence, the sources in Fig. A4 match with the left halves of the sources in Fig. A2 and the dispersal distance is underestimated for the Gaussian kernel but overestimated for the power-law kernel.

For the power-law kernel, the kernel parameter is substantially underestimated across the whole range of parameters: green curves in Fig. A3A, A3B correspond to negative errors. For this reason, the kernel parameter estimates are in the range γ < 3 (while the true values are in the range γ > 3), leading to an infinite mean dispersal distance, 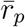 (this is a mathematical property of power-law kernels; see Box 1).

**Figure A3:**
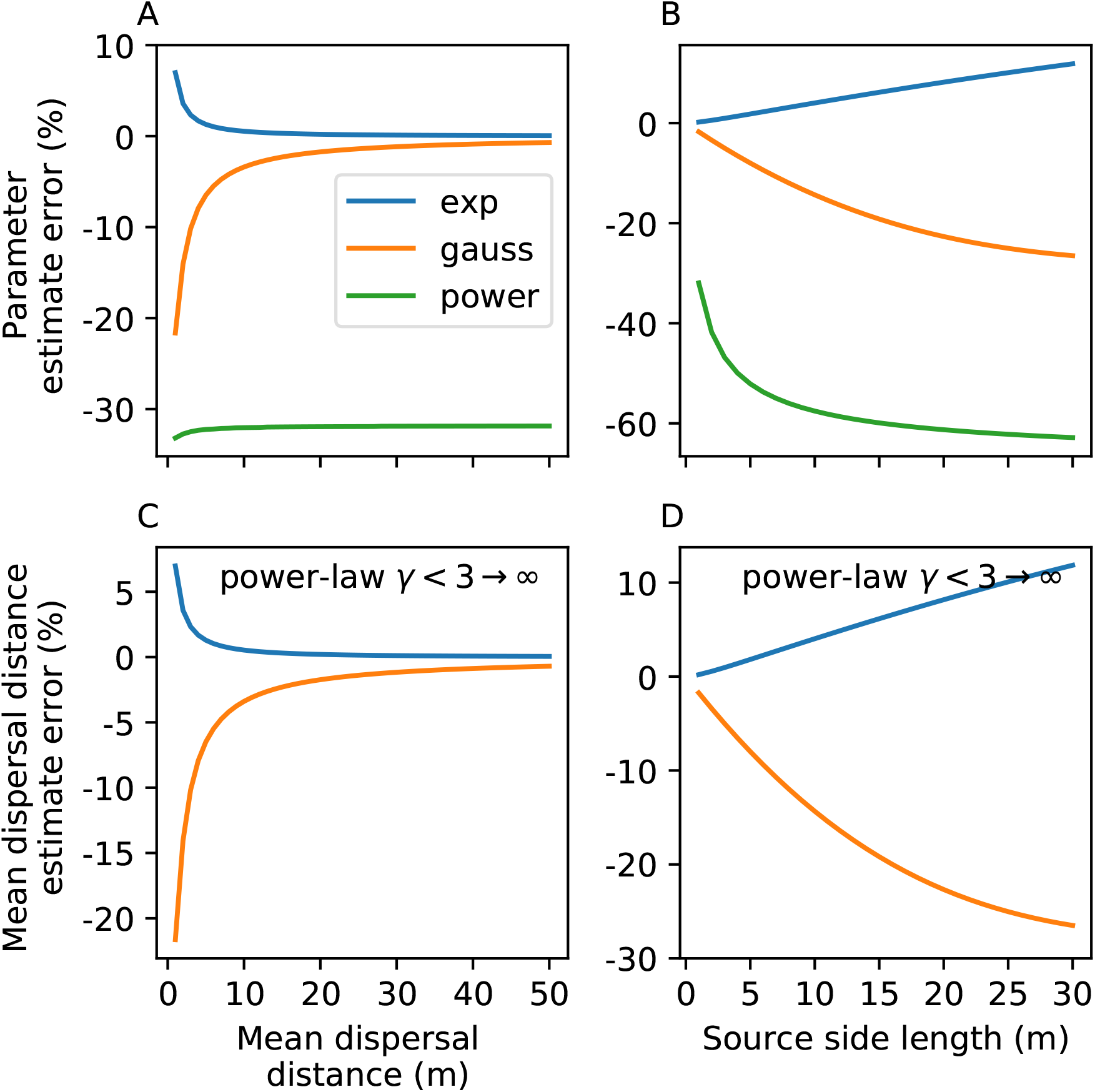
The accuracy of the point-source approximation depends on the true values of mean dispersal distances (left column) and on the source size (right column). Simulation design is the same as in Fig. A1, the only difference is that here a virtual point source is at the edge of the square source area, not at the center. Parameter estimates become more accurate when considering organisms with longer mean dispersal distances and when using smaller sources. Parameters: see Fig. A1.

**Figure A4:**
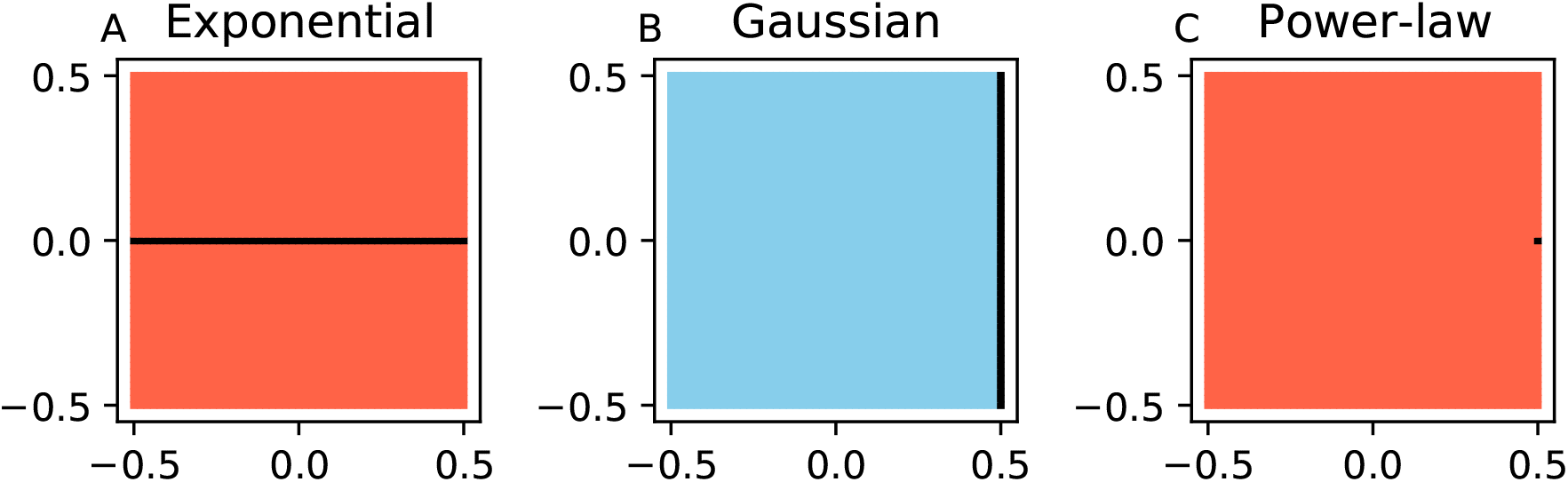
Contributions of individual points within the source area to the errors in the estimates of mean dispersal distances, 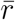. Here, we considered a virtual point source located at the edge of the actual square source area. For each point within the source area (across a 2 cm × 2 cm grid) the color shows whether the point contributes to an underestimation (blue), or an overestimation (red) of the mean dispersal distance, or represents an accurate estimate (black). Measurements are conducted on the right side of the source area. Parameters: see Fig. A2.

We conclude that it is difficult to guess *a priori* how the accuracy of estimation of the kernel parameters and mean dispersal distances would be affected by specific extensions of the source area or positioning of the virtual point source within the source area. Therefore, we are not able to provide a simple rule of thumb regarding the validity of the point-source approximation. Thus, using the spatially explicit approach is recommended to achieve more accurate estimation of dispersal kernel parameters.

### Estimation of power-law kernel parameters is problematic under the point-source approximation

In Figure 5A in the main text, the error of the kernel parameter estimate approached zero quite fast for exponential and Gaussian kernels as we increased the mean dispersal distance. In contrast, for the power-law kernel, after an initial decline the error approached a constant value of around 3% at long mean dispersal distances. We wanted to better understand the reasons behind this qualitative difference and to investigate whether we could further reduce this remaining error. For this purpose, we formulated three hypotheses and tested them by modifying the simulation and fitting procedures. Each hypothesis assumes a specific reason behind the impaired accuracy of estimation of the power-law kernel parameter: (i) insufficiently dense sampling of the dispersal gradient (i.e., insufficient number of measurement lines within the fixed length of the measured gradient); (ii) insufficient capturing of the tail of the dispersal gradient; and (iii) fitting only the shape parameter γ of the power-law kernel, while setting the scale parameter λ to a constant value.

**Figure A5:**
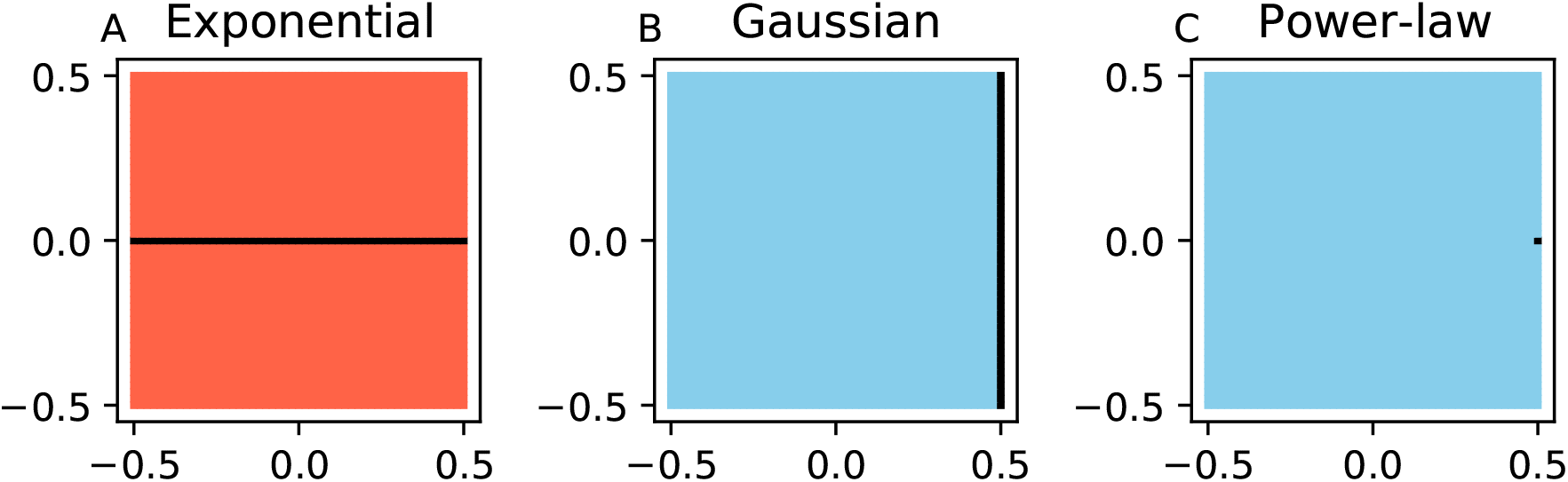
Contributions of individual points within the source area to the errors in the kernel parameter estimates. Here, we considered a virtual point source located at the edge of the actual square source area (in contrast to Fig. 6 in the main text that presents analogous outcomes when the virtual point source is at the center of the source area). For each point within the source area (across a 2 cm × 2 cm grid) the color shows whether the point contributes to an underestimation (blue), or an overestimation (red) of the kernel parameter, or represents an accurate estimate (black). Measurements are conducted on the right side of the source area. Parameters: see Fig. A2.

To test the first hypotheses, we increased the density of measurement lines. To test the second hypothesis, we introduced additional measurements further away from the source. To test the third hypothesis, we set the scale parameter of the power-law kernel (λ) free to be estimated alongside with the shape parameter γ. In these simulations, we used a 1 m-by-1 m source and set the mean dispersal distance to 20 m as in Fig. 5.

Before introducing these modifications, we re-state the “baseline” results. As explained in the main text, we used the 10 cm distance between adjacent measurement lines and continued sampling until we reached 1% of the intensity in the first measurement line. In this case, the kernel parameter estimate was γ_es_t = 3.153. This estimate exhibits a 3.37% overestimation compared to the true value γ_true_ = 3.05. The associated mean dispersal distance was estimated to be 6.54 m, a severe underestimation of the true value of 20 m. (Note how a small error in the parameter estimate translates into a substantial error in the estimate of the mean dispersal distance when γ is close to its critical value of 3.)

#### (i) Sample more densely

We reduced the distance between adjacent measurement lines from 10 cm to 1 cm, thereby introducing a ten-fold increase in the density of measurements. We also started measurements from 1 cm away from the edge of the source, instead of 10 cm originally. As a result, the error in estimation decreased slightly from 3.37% to 2.25% overestimation (γ_est_ = 3.119) and the estimated mean dispersal distance was now 8.4 m, also slightly closer to the true value of 20 m.

#### (ii) Sample further away from the source

We decreased the threshold to stop the sampling: now we included all points that had at least 0.1% of the intensity of the first sampling line (the original threshold was 1%). This led to the estimate *γ_est_* = 3.152 (an overestimation of 3.36%) corresponding to 6.56 m estimated mean dispersal distance. This represented only a negligible improvement in the estimation accuracy.

#### (iii) Estimate A parameter together with γ-parameter

We fitted the parameter λ together with the parameter γ. This improved quality of the 1D fit, leading to the chi-squared value of 0.699 compared to 1.278 in the original fit. However, the error in the shape parameter estimate increased (γ_*est*_ = 3.337 (error: 9.33%)) and so did the error in the estimate of the mean dispersal distance (3.50 m; *λ_est_* = 0.585).

### Conclusion

We introduced specific modifications of the sampling and fitting procedures to test the three hypotheses described above. In each of the three cases, we achieved relatively minor improvements either in the accuracy of parameter estimates or in the quality of fits. The largest improvement in the estimation accuracy was gained by the ten-fold increase in sampling density: this reduced the error from 3.37% to 2.25%. However, this improvement does not appear satisfactory considering the increased cost of denser sampling. When sampling further away from the source (thereby capturing regions further into the tail of the dispersal gradient), the improvement in the estimation accuracy was negligible, while the practical challenges in detecting dispersing populations at 0.1% of their highest density would likely be considerable. Finally, fitting both parameters simultaneously improved the quality of the fit, but impaired the accuracy of parameter estimation. Thus, the one-dimensional fitting under the point-source approximation has serious limitations for parameter estimation of power-law dispersal kernels. These limitations can be overcome by using the spatially explicit method: it provided more accurate estimates in every case of our simulated dispersal experiments.

## Appendix B: Realistic features of experimental design influence the estimation accuracy of the power-law kernel parameter under the point-source approximation

In the “Simulations” section of the main text, we considered a “more realistic scenario”, where we simulated measurements of dispersal in a plausible experimental design. We accounted for the fact that when measuring dispersal gradients, each sampling location is distributed over a finite area and sampling is conducted a limited number of times. In this scenario, the kernel parameter estimates were not entirely accurate even when using the spatially explicit approach. Moreover, even in the idealized simulations, the error of the parameter estimate did not approach zero with increasing mean dispersal distances for the power-law kernel (green curve in Fig. 5A). We wanted to understand the reasons behind those remaining errors. Hence, here we present more detailed, additional simulations we conducted using the power-law kernel. We investigated modifications to the “realistic” scenario, showing that some of the inaccuracies can be countered by changes in the experimental design. Results of similar simulations are available for exponential and Gaussian kernels in the data repository (Data dryad, TBA).

~~~
Data: https://www.dropbox.com/t/YE3oj8Bo73uiEziE
~~~

We conducted additional simulations with slight modifications with respect to the ones described in subsection “Estimation of dispersal kernel parameters in a more realistic scenario” of the “Simulations” section of the main text. In the main text, measurement lines were 40m-long and were sampled at five points within each line. Here, we used measurements lines of four different lengths (1 m, 5 m, 20 m and 40 m) and varied the number of sampling points within the measurement lines from 5 to 10, 40 and 160.

### Effect of sampling density on the estimation accuracy of the power-law kernel parameter

The inputs to the fitting procedure were means over all measurements within each measurement line, and the fitting assumed dense, uniform sampling (simulation design in Fig. 4C). Hence, we hypothesized that higher sampling density within measurement lines would improve the accuracy of estimation. We were also interested in the sampling density required to achieve a desired level of accuracy.

We found that more dense sampling (achieved by increasing the number of sampling points while maintaining the size of the measurement lines) indeed led to more accurate estimates in both the spatially explicit approach (2D-estimation) and under the point-source approximation (1D-estimation). Importantly, at an increased number of sampling points, the 2D-estimates did approach their true value, while the 1D-estimates retained substantial errors (Tables B1 and B2).

### Effect of the spatial extension of destinations on the estimation accuracy of the power-law kernel parameter

We hypothesized that the length of measurement lines (brown vertical lines in Fig. 4C) introduced error in the estimation, as the number of dispersed individuals measured at the ends of the measurement lines (orange lines in Fig. 4C) would generally be different compared to the one measured in the middle of the lines. We conducted simulations with shorter measurement lines to test this hypothesis and to determine whether shorter measurement lines would result in more accurate estimates.

We found that the estimates with shorter measurement lines were generally more accurate (Tables B1 and B2), thereby confirming our hypothesis. As we shortened the measurement lines, 2D-estimates approached their true value, while 1D-estimates retained substantial errors.

**Table B1:** Estimation of the power-law kernel parameter after simulating dispersal from a square source of 10 m × 10 m, with variable sampling density.

| Measurement lines (m) | # of sampling points | 1D | 2D | True |
| --- | --- | --- | --- | --- |
| 1 | 5 | 5.06 | 5.00 | 5.0 |
| 1 | 10 | 5.06 | 5.00 | 5.0 |
| 1 | 40 | 5.06 | 5.00 | 5.0 |
| 1 | 160 | 5.06 | 5.00 | 5.0 |
| 5 | 5 | 5.03 | 4.99 | 5.0 |
| 5 | 10 | 5.04 | 5.00 | 5.0 |
| 5 | 40 | 5.04 | 5.00 | 5.0 |
| 5 | 160 | 5.04 | 5.00 | 5.0 |
| 20 | 5 | 4.72 | 4.90 | 5.0 |
| 20 | 10 | 4.78 | 4.96 | 5.0 |
| 20 | 40 | 4.81 | 4.99 | 5.0 |
| 20 | 160 | 4.82 | 5.00 | 5.0 |
| 40 | 5 | 4.27 | 4.83 | 5.0 |
| 40 | 10 | 4.35 | 4.92 | 5.0 |
| 40 | 40 | 4.41 | 4.98 | 5.0 |
| 40 | 160 | 4.42 | 5.00 | 5.0 |

Note: “2D” and “1D” stand for two- and one-dimensional models, respectively. The scale parameter of the power-law kernel is set to $\lambda = 20 \text{ m}$ .

Furthermore, we noticed that errors in the estimates of kernel parameters caused by the point source approximation in the more realistic scenario (Table 2, column “1D”) were consistent with the errors caused by the point-source approximation in the idealized simulations (where the destinations were considered as points) shown in Fig. 5 in the case of exponential and Gaussian kernels, but not in the case of the power-law kernel. For exponential and Gaussian kernels, the kernel parameters were overestimated both in the idealized simulations (Fig. 5) and in more realistic simulations Table 2 (column “1D”; more realistic simulations). In contrast, for the power-law kernel, the kernel parameter was overestimated in the idealized simulations, but underestimated in more realistic simulations. We wanted to find out why.

For this purpose, we focused on the subset of simulations with square source and five sampling points in each measurement line (these are extracted from Table B1 into Table B3). The kernel parameter was underestimated with 1 m and 5 m measurement lines (in agreement with idealized simulations) but overestimated with longer lines (column “1D” in Table B3). Hence, the extension of the destinations under the point-source approximation affected not only the magnitude of the estimate error but also its sign. This may explain why the sign of errors differed between the idealized and the more realistic simulations.

**Table B2:** Estimation of the power-law kernel parameter after simulating dispersal from a circle source of 20 m diameter, with variable sampling density.

| Measurement lines (m) | # of sampling points | 1D | 2D | True |
| --- | --- | --- | --- | --- |
| 1 | 5 | 5.20 | 5.00 | 5.0 |
| 1 | 10 | 5.20 | 5.00 | 5.0 |
| 1 | 40 | 5.20 | 5.00 | 5.0 |
| 1 | 160 | 5.20 | 5.00 | 5.0 |
| 5 | 5 | 5.19 | 4.99 | 5.0 |
| 5 | 10 | 5.19 | 5.00 | 5.0 |
| 5 | 40 | 5.19 | 5.00 | 5.0 |
| 5 | 160 | 5.19 | 5.00 | 5.0 |
| 20 | 5 | 4.95 | 4.93 | 5.0 |
| 20 | 10 | 4.99 | 4.97 | 5.0 |
| 20 | 40 | 5.02 | 4.99 | 5.0 |
| 20 | 160 | 5.02 | 5.00 | 5.0 |
| 40 | 5 | 4.58 | 4.85 | 5.0 |
| 40 | 10 | 4.62 | 4.93 | 5.0 |
| 40 | 40 | 4.67 | 4.98 | 5.0 |
| 40 | 160 | 4.68 | 5.00 | 5.0 |

Note: “2D” and “1D” stand for two- and one-dimensional models, respectively. The scale parameter of the power-law kernel is set to $\lambda = 20$ m.

A similar change in the sign of the estimation error is also observed with circular source areas. Interestingly, in certain cases with circular source, elongated measurement lines and sparse sampling, the 1D-estimation resulted in more accurate estimates than the 2D-estimation. Note that the 1D-estimate happens to be more accurate than the 2D-estimate with 20 m measurement lines and 5 or 10 sampling points per line (column “1D” in Table B2). In these cases, the estimation errors due to the extension of the source opposed the effect of the extension of measurement lines, which “accidentally” led to more accurate estimates since the two errors compensated for each other.

**Table B3:** Estimation of the power-law kernel parameter after simulating dispersal from a square source of 10 m × 10 m, varying the length of measurement lines (with five sampling points).

| Measurement lines (m) | 1D | 2D | True |
| --- | --- | --- | --- |
| 1 | 5.06 | 5.00 | 5.0 |
| 5 | 5.03 | 4.99 | 5.0 |
| 20 | 4.72 | 4.90 | 5.0 |
| 40 | 4.27 | 4.83 | 5.0 |

Note: “2D” and “1D” stand for two- and one-dimensional models, respectively. The scale parameter of the power-law kernel is set to $\lambda = 20\text{ m}$ .

Considered together, these outcomes highlight the importance of using appropriate, spatially explicit modelling and simulations when designing dispersal experiments and analyzing data.

